# Monitoring GPCR Conformation with GFP-Inspired Dyes

**DOI:** 10.1101/2023.06.01.543196

**Authors:** Anatoliy Belousov, Ivan Maslov, Philipp Orekhov, Polina Khorn, Pavel Kuzmichev, Nadezhda Baleeva, Vladislav Motov, Andrey Bogorodskiy, Svetlana Krasnova, Konstantin Mineev, Dmitry Zinchenko, Evgeni Zernii, Valentin Ivanovich, Sergei Permyakov, Johan Hofkens, Jelle Hendrix, Vadim Cherezov, Thomas Gensch, Alexander Mishin, Mikhail Baranov, Alexey Mishin, Valentin Borshchevskiy

**Affiliations:** Moscow Institute of Physics and Technology, Dolgoprudny, 141701, Russia; Dynamic Bioimaging Lab, Advanced Optical Microscopy Centre, Biomedical Research Institute, Agoralaan C (BIOMED), Hasselt University, Diepenbeek, 3590, Belgium; Laboratory for Photochemistry and Spectroscopy, Division for Molecular Imaging and Photonics, Department of Chemistry, KU Leuven, Leuven, 3001, Belgium; Faculty of Biology, Shenzhen MSU-BIT University, Shenzhen, 518172, China; Institute of Bioorganic Chemistry, Russian Academy of Sciences, Moscow, 117997, Russia; Pirogov Russian National Research Medical University, Moscow, 117997, Russia; National Research University Higher School of Economics, Moscow, 101000, Russia; Branch of Shemyakin and Ovchinnikov Institute of Bioorganic Chemistry, Russian Academy of Sciences, Pushchino 142290, Russia; Belozersky Institute of Physico-Chemical Biology, Lomonosov Moscow State University, Moscow, 119992, Russia; Institute for Biological Instrumentation, Russian Academy of Sciences, Pushchino, 142292, Russia; Max Plank Institute for Polymer Research, Mainz, Germany; Department of Chemistry, University of Southern California, Los Angeles, CA 90089, USA; Joint Institute for Nuclear Research, Dubna 141980, Russian Federation; Current affiliation: Goethe-University Frankfurt, Frankfurt, Germany

## Abstract

Solvatochromic compounds have emerged as valuable environment-sensitive probes for biological research, with the chromophore of the green fluorescent protein (GFP) being a well-studied example. In this study, we demonstrate that synthetic analogues of the GFP chromophore can be used to investigate ligand-induced conformational changes in proteins. We synthesized thiol-reactive derivatives of four analogues of the GFP chromophore that exhibit notable solvatochromism. We used these derivatives to label two proteins: the soluble calcium sensor recoverin (Rec) and the transmembrane G protein-coupled A_2A_ adenosine receptor (A_2A_AR), via cysteines located or introduced in the regions that undergo structural changes upon ligand binding. Two of these dyes showed Ca^2+^-induced fluorescence changes when attached to Rec. Notably, our best-performing dye, DyeC, when attached to A_2A_AR, revealed agonist-induced changes in both fluorescence intensity and shape of the emission spectrum. Molecular dynamics (MD) simulations provided mechanistic insights into these changes showing the activation of A_2A_AR transfers DyeC to a more confined and more hydrophilic environment. Additionally, an allosteric modulator, HMA, induces changes in DyeC fluorescence spectra, indicating a distinct receptor conformation from apo, antagonist, or agonist-bound receptors. Our study demonstrates that GFP-inspired dyes are effective for detecting structural changes in GPCR (G protein-coupled receptors), with advantages such as the ability to perform both intensity-based and ratiometric tracking, red-shifted fluorescence spectra, high extinction coefficient, and sensitivity to allosteric modulation. These dyes expand the toolbox for tracking ligand-induced changes and facilitate new insights into conformational changes induced by allosteric modulators in GPCRs.

## Introduction

From in vivo imaging to single-molecule tracking, the green fluorescent protein (GFP) has become an indispensable tool for many biological studies [1]. The GFP chromophore, 4-hydroxybenzylidene-dimethylimidazolinone (HBDI, Figure 1), is spontaneously formed due to specific cyclization of three amino acid residues located in the center of the GFP β-barrel. Structurally modified synthetic analogues of the GFP chromophore represent a diverse class of benzylidene imidazolones (BDI) that found many applications as versatile labels due to their exceptional fluorescent properties, small size, and easy synthesis [2]. Similar to GFP, where the high fluorescence yield of HBDI is supported by the protein microenvironment, the transition of some BDI derivatives from water to less polar solvents results in a substantial increase in their fluorescence quantum yield (FQY). Such compounds have been extensively tested as fluorogenic environmentally sensitive dyes [3–5].

**Figure 1.**
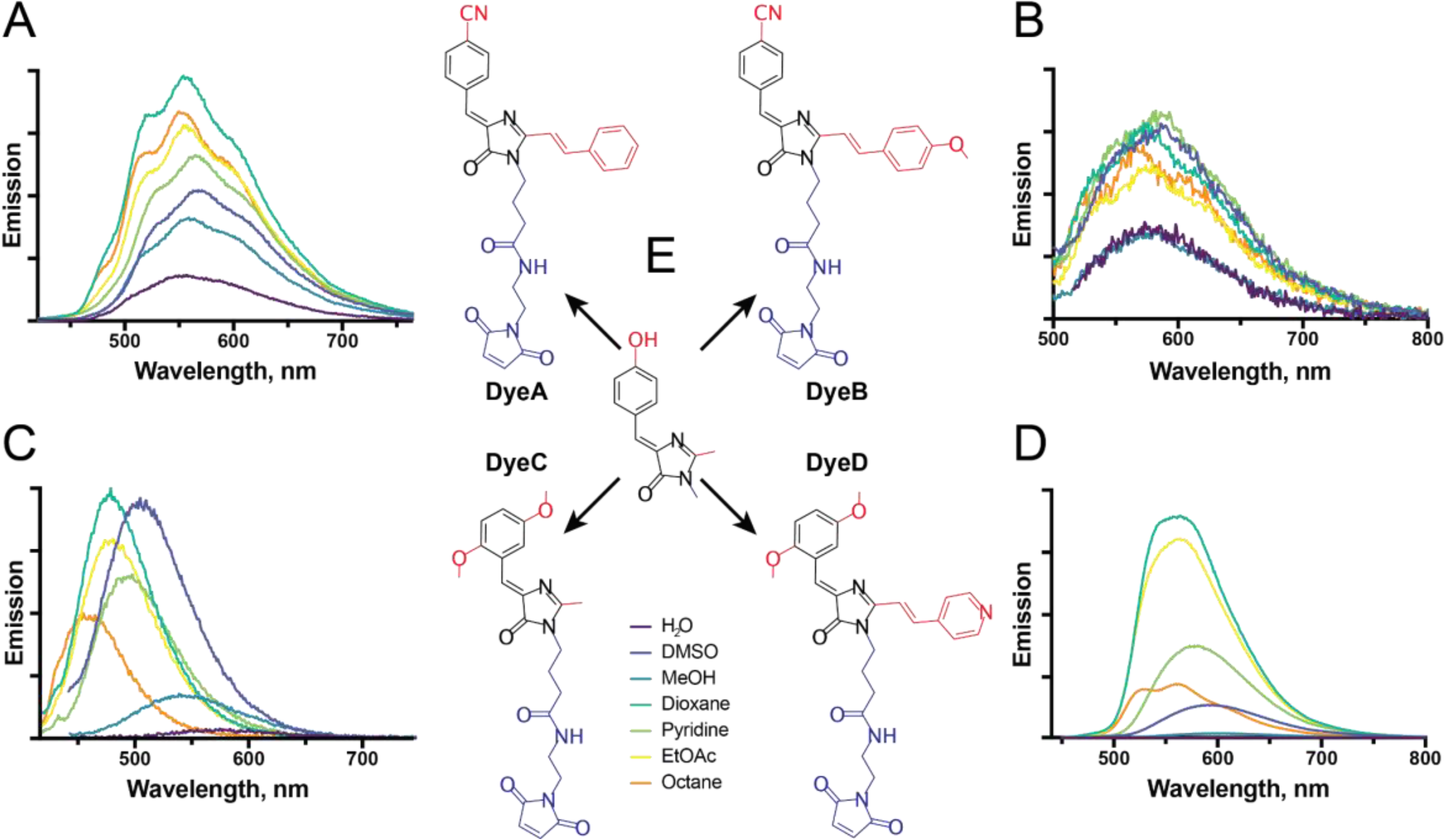
Dye structures and their emission spectra in various solvents. **A-D**: Changes of the emission fluorescence spectra for free DyeA **(A),** DyeB **(B)**, DyeC **(C)**, and DyeD **(D)** in solvents with different polarity and viscosity. The excitation wavelengths were 410 nm (DyeA), 430 nm (DyeB), 380 nm (DyeC), and 420 nm (DyeD). Complete results and additional information for the tested dyes are provided in Supplementary Table S1. **E:** The HBDI fluorescent core is shown in the center and four dyes as its derivatives; the maleimide groups, responsible for cysteine interaction, are colored in blue; the fluorescent core modifications are colored in red.

Environmentally sensitive labels have often been used to study ligand-induced structural changes in proteins [6–12]. Typically, in this approach, their thiol-reactive variants are attached to cysteine residues (either intentionally introduced or naturally present) located in the vicinity of functionally labile protein elements. In successful constructs, fluorescence properties of the label have the ability to report on protein activation or inhibition. These studies shed light on the molecular mechanisms of protein activation and in some cases facilitated the creation of biomolecular sensors for a wide variety of metabolites [8,13–18]. Despite the wide range of applications, GFP-inspired labels have never been used for this purpose before. In this study, we show these labels can serve as environmentally sensitive labels to report on conformational changes in GPCRs.

GPCRs constitute the largest class of membrane proteins in humans that regulate critical physiological processes, e.g. vision, taste, neurotransmission, and inflammation. More than one third of drugs approved by the United States Food and Drug Administration (FDA) have GPCRs as their primary targets [19]. Most of these drugs target the primary extracellular binding site that naturally accommodates endogenous ligands, i.e. the orthosteric ligand-binding pocket. The ligand binding causes structural changes propagating across the receptor towards the intracellular side and enables coupling to its cognate G-proteins or other intracellular partners. In the absence of ligands, most GPCRs exhibit a certain level of activity known as basal signaling. Orthosteric ligands can directly control GPCR activity: agonists increase the basal signaling, antagonists occupy the ligand-binding site but do not affect the receptor’s activity, and inverse agonists decrease the basal signaling [20]. The affinity and efficacy of orthosteric ligands can be affected by allosteric modulators that bind to spatially distinct sites on GPCRs and modulate their function [21–24]. Notably, some allosteric modulators can affect receptor’s activation on their own [23–27].

The A_2A_ adenosine receptor (A_2A_AR), one of the most studied GPCRs, is a promising target for drugs against cancer, chronic pain, sleep disorders, depression, and Parkinson’s disease [28]. It regulates cardiovascular system and inflammation throughout the body and modulates the neurotransmission of glutamate and dopamine in the brain [29,30]. A_2A_AR has often been used as a prototypical receptor for the development of biophysical techniques for studying other GPCRs [31–35].

Here we assessed four GFP-inspired fluorophores for their potential to serve as environmentally sensitive labels to report on conformational changes in proteins. For this, we attached a maleimide group to them for cysteine labeling and studied their spectral properties in solvents with varying polarity and viscosity. The best of them were then employed to label two proteins: bovine recoverin (Rec) and A_2A_AR. We used Rec as a convenient model protein to evaluate the performance of environmentally sensitive labels. This is attributed to the presence of a sole cysteine residue and the considerable conformational alterations experienced upon activation by Ca^2+^. Using the best performing dye attached to A_2A_AR, we investigated effects of various ligands, including antagonists, agonists, and an allosteric modulator, on its fluorescence and observed reliable and distinctive changes. Finally, we conducted molecular dynamic (MD) simulations of the A_2A_AR complexes with the dyes to obtain structural insights into the observed changes.

## Results

### Synthesized thiol-reactive GFP-inspired fluorophores show solvatochromism

To follow conformational changes in proteins using GFP-inspired fluorophores, we selected BDI derivatives that previously showed solvatochromic properties. It has been reported that the FQY of 4-nitrile and 2,5-dime-thoxy derivatives of a BDI benzyli-dene fragment with the absorption maxima in the UV range depends on solvent [36,37]. Since fluorophores with red-shifted absorption spectra are preferred in many quantitative fluorescence-based measurements due to their lower phototoxicity, background autofluorescence, and scattering, we used their BDI imidazolone derivatives, which have shown a red-shifted absorption together with a remarkable solvatochromism of their emission maxima and high FQY [3]. Therefore, we selected two 4-nitrile (Figure 1, DyeA and DyeB) and two 2,5-dime-thoxy (Figure 1, DyeC and DyeD) benzyli-dene derivatives of red-shifted BDI compounds and complemented them with a maleimide group to enable covalent binding to cysteine residues.

To assess the solvatochromism of the maleimide-conjugated compounds, we investigated their fluorescence properties in solvents with different polarities ranging from water to pentadecane (Figure 1, Supplementary Table S1). Similarly to the original compounds, all four dyes showed notable changes in their fluorescence emission spectra depending on the solvent. We observed that FQYs of the 4-nitrile benzyli-dene compounds (DyeA and DyeB) strongly depend on the solvent polarity, while the wavelengths of their emission maxima vary by <10 nm in different solvents. At the same time, the 2,5-dime-thoxy benzyli-dene compounds (DyeC and DyeD) showed solvent-dependent changes both in FQY and the wavelength of their emission maxima.

### Fluorescence of GFP-inspired dyes highlights Ca^2+^-dependent conformational **changes in Rec**

To select dyes suitable for tracking ligand-induced structural changes in proteins, we used a water soluble protein Rec. Rec is a 23 kDa member of the neuronal calcium sensor family [38], which participates in regulation of phototransduction and light adaptation of retinal photoreceptors [39]. Rec has two calcium binding sites, and the interaction with calcium ions results in substantial changes in its structure [40]. Rec contains a single cysteine residue, which is conveniently located in the protein loop subjected to structural changes upon activation [41]. The emission spectrum of the Alexa647 label bound to this cysteine was shown to undergo substantial changes depending on the calcium concentration [8]. All these results and considerations make Rec a perfect test object to validate whether our modified BDI fluorescent labels can report structural changes in a protein.

We labeled Rec with each of the four synthesized maleimide dyes. The absorption spectra of labeled Rec (Supplementary Figure S1) showed high labeling efficiencies for DyeA, DyeB, and DyeC (∼90–100%), while DyeD had a low labeling efficiency (<5%) and, therefore, was excluded from further experiments. The low labeling efficiency is probably related to the low water solubility of DyeD that was originally observed for its parent compound (5d in [3]).

We measured fluorescence emission spectra of the labeled Rec in Ca^2+^-bound and Ca^2+^– free states (Figure 2). Rec-DyeB and Rec-DyeC displayed pronounced Ca^2+^-induced changes in their fluorescence spectra. Similarly to previously characterized solvatochromic properties of DyeB, Rec-DyeB showed a higher emission intensity in the presence of calcium; however, its emission maximum also red-shifted by >50 nm, while the emission wavelength of free DyeB did not exhibit a solvatochromic behavior. On the other hand, Rec-DyeC showed a 50 nm red-shift of the emission maximum as expected from the solvatochromic behavior of the free dye, but without any changes in the fluorescence intensity. We further tested the labeling specificity for our best-performing dye, DyeC, and observed that the efficiency of non-specific labeling of the cysteine-free Rec mutant Rec_C39D_ was negligible (<5%, Supplementary Figure S1), suggesting that C39 was the only labeled amino acid in the WT protein.

**Figure 2.**
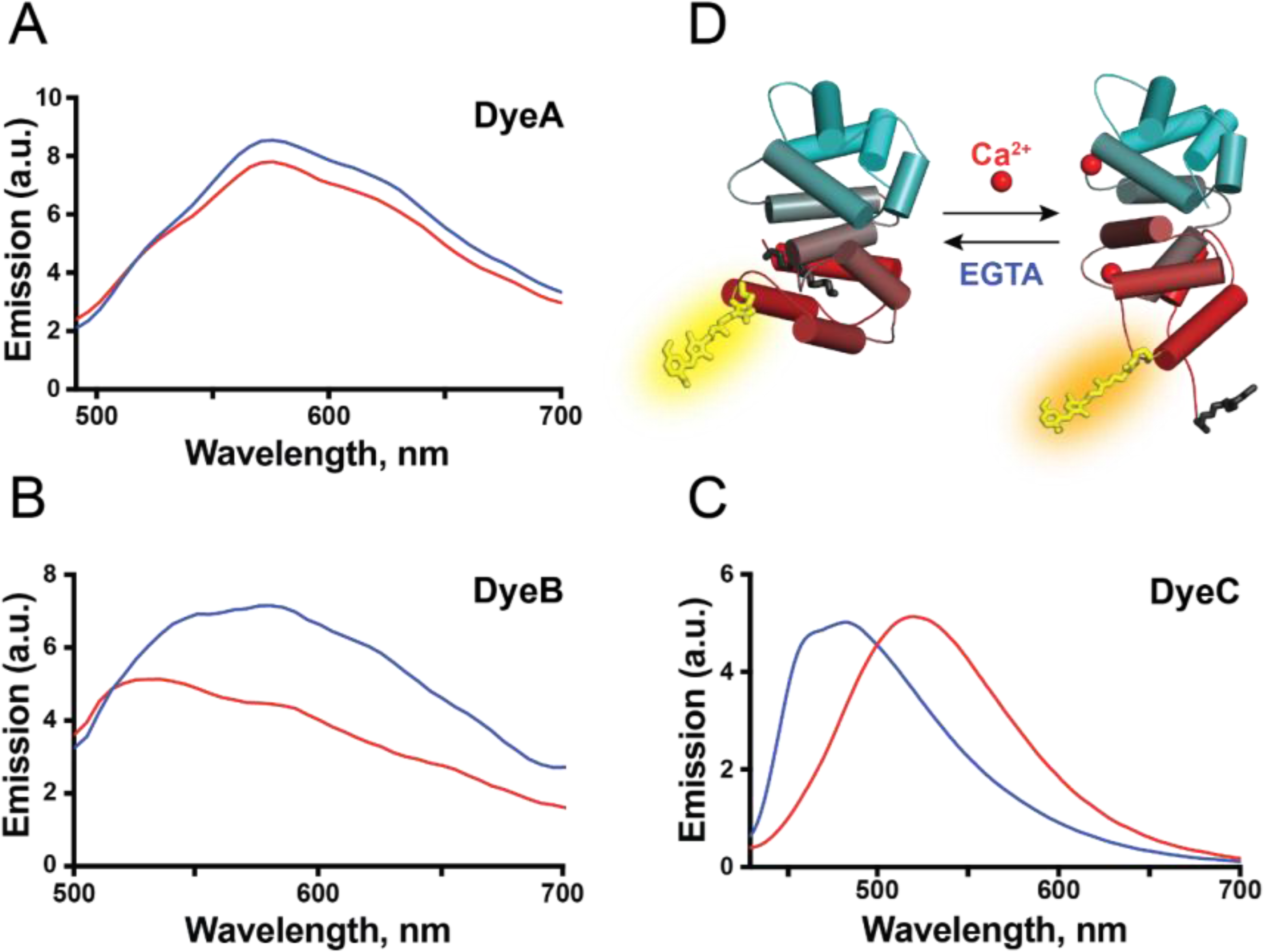
Ca^2+^-induced conformational and spectral changes of Rec labeled with DyeA, DyeB, and DyeC. **A-C:** The Ca^2+^-induced response of Rec with three BDI derived labels (DyeA, DyeB, and DyeC). Blue curves correspond to samples in the presence of 100 µM CaCl2, red curves correspond to calcium-free samples with 100 µM of chelator EGTA. The protein concentration was maintained at 10 µM. Excitation wavelengths were 440 nm, 460 nm, and 410 nm for DyeA, DyeB and DyeC, respectively. **D:** Structural rearrangements of labeled Rec induced by calcium ions. The structures of the calcium-free and calcium-bound forms are based on 1IKU and 1JSA PDBs, respectively [40,82]. DyeC attached to the single native cysteine (C39) in Rec is shown in yellow, the myristoyl group at the N-terminus of Rec is shown in black. Rec is colored in gradient from red on its N-terminus to cyan on C-terminus.

### Fluorescence of DyeC highlights conformational changes in A2AAR induced by agonists and an allosteric modulator

To track ligand-induced conformational changes in a GPCR, we labeled A_2A_AR with DyeB and DyeC. For this purpose, we applied a single-point mutation L225^6.27^C (superscripts indicate Ballesteros–Weinstein numbering [42]) to introduce a cysteine residue at the intracellular tip of the transmembrane helix TM6. A large-scale movement of the intracellular part of TM6 is one of the hallmarks of activation in class A GPCRs, including A_2A_AR [43]. Similar label placements at the intracellular tip of TM6 were used in previous fluorescence-based [44,45] and F^19^-NMR [32,46,47] studies.

The wild-type A_2A_AR (A_2A_AR_WT_) has 6 native unpaired cysteines buried in the receptor core. To attach dyes only to the introduced cysteine, we labeled A_2A_AR_L225C_ in crude membranes as described previously [45,48]. After labeling, A_2A_AR_L225C_ was purified and reconstituted in lipid nanodiscs (ND). The labeling efficiency was estimated as 80% for A_2A_AR_L225C_-DyeC and <5% for A_2A_AR_L225C_-DyeB (Supplementary Figure S2), therefore, DyeB was excluded from further experiments. As a negative control for labeling specificity, we also tried labeling A_2A_AR_WT_ with DyeC. The A_2A_AR_WT_-DyeC sample showed a very low but detectable fluorescence signal indicating a small percentage of non-specific labeling (<10%, Supplementary Figure S2). However, the fluorescence emission spectrum of A_2A_AR_WT_-DyeC did not change upon the addition of A_2A_AR ligands (Supplementary Figure S3), and, therefore, we concluded that the non-specifically labeled A_2A_AR did not contribute to the observed spectral changes described below.

To test the response of DyeC to A_2A_AR activation, we measured its fluorescence emission spectra (Figure 3B) in the apo state, as well as in complex with two agonists (NECA and adenosine) and two antagonists (ZM241385 and SCH58261). The A_2A_AR-DyeC emission spectra showed a similar response to both agonists, while the antagonists did not change the emission spectra compared to the apo state. The agonists increased the integrated fluorescence intensity by ∼20%, which we quantified as the integrated fluorescence intensity of the emission spectrum (Figure 3C). Furthermore, the agonists induced a red shift of ∼5 nm in the emission maximum and resulted in a steeper left shoulder in the spectrum. To quantify this change, we used the ratio between the intensities of the main emission peak at 520 nm and the blue-shifted shoulder peak at 460 nm (I_520_/I_460_). The ratio I_520_/I_460_ changed from ∼1.7 for the apo and antagonist-bound receptor to ∼2.0 for agonist-bound receptor (Supplementary Table S2). Thus, DyeC allowed tracking of the conformation changes in A_2A_AR based on changes in both the fluorescence intensity and the shape of the spectrum.

**Figure 3.**
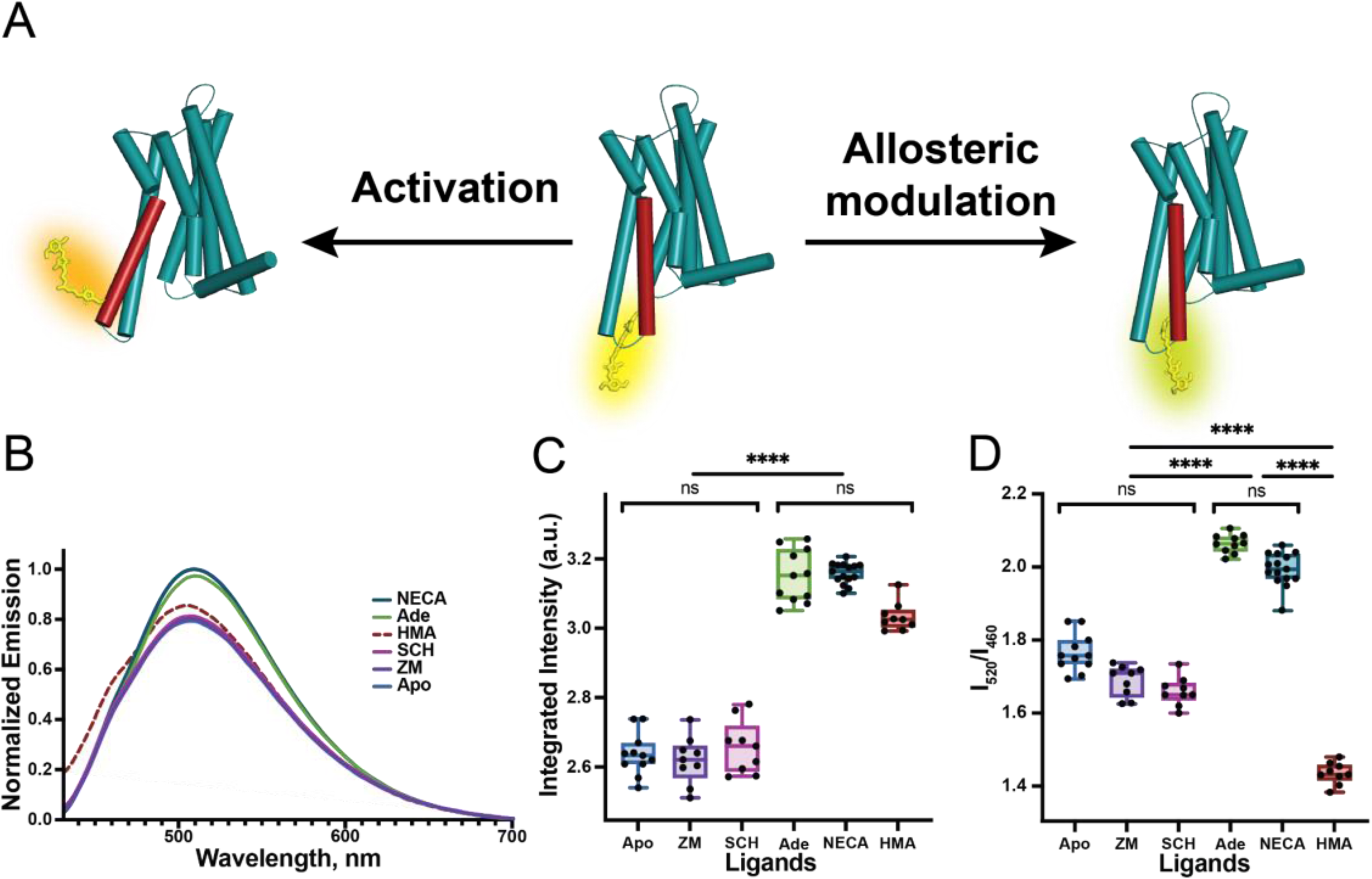
Structural and spectral changes of A2AARL225C-DyeC induced by various ligands. **A:** Schematic representation of structural changes caused by agonists and the allosteric modulator HMA in A2AARL225C-DyeC. Structures of the active and non-active A2AAR are sketched from 6GDG and 3RFM, respectively [34,70]. Agonist binding results in an outward shift of the intracellular part of the TM6. The structural effects of the allosteric modulator HMA remain unknown. **B:** Emission spectra of A2AARL225C-DyeC bound to antagonists (ZM241385 and SCH58261), agonists (NECA and Adenosine), allosteric modulator (HMA), and in apo state. **C:** Variation of the integrated intensity of A2AARL225C-DyeC bound to different ligands. The integrated intensity is quantified as the area under the fluorescence emission spectrum from 430 to 700 nm. **D:** Variation of the intensity ratio I520/I460 for different ligands. Each condition in **C** and **D** was measured at least 9 times with protein from at least three independent purifications, each protein sample was mixed with the ligand independently. The data represent the mean±SD. The protein concentration was maintained at 10 µM, all ligands were added at saturating concentration of 100 µM. The significance level is given according to the ordinary one-way ANOVA with the post hoc Tukey HSD test: ∗∗p < 0.005, ∗p < 0.05, ns – not significant.

To test whether ligand-induced changes in A_2A_AR_L225C_-DyeC fluorescence spectra were due to conformational changes of the receptor and not due to direct interaction between DyeC and ligands, we conducted displacement experiments of agonist NECA with the excess of antagonist ZM241385, and viсe versa (Supplementary Figure S4). NECA and ZM241385 are both highly specific orthosteric A_2A_AR ligands (with nanomolar Kd’s [https://www.guidetopharmacology.org]). The fact that the effect of NECA can be fully reversed by an excess of ZM241385, and vice versa, indicates that both ligands can displace each other from the orthosteric site and have no direct interactions with DyeC.

After establishing the response to orthosteric agonists and antagonists, we used A_2A_AR_L225C_-DyeC to characterize structural changes in A_2A_AR induced by one of the most potent allosteric ligands, 5-(N,N-hexamethylene)-amiloride (HMA, Figure 3B). Previous docking simulations suggested that HMA binds to a conserved intramembrane sodium-binding site that is spatially distinct from the orthosteric binding site in A_2A_AR, where all other ligands used in this study bind [49]. Previous functional studies showed that HMA increases the dissociation rate (*k*_off_) for A_2A_AR antagonists, but not agonists [50,51]. At the same time, HMA was shown to displace both antagonists and agonists in competition assays [49,50,52]. In our experiments, the shape of the fluorescence emission spectrum for the A_2A_AR_L225C_-DyeC complex with HMA differed significantly from those for the receptor bound to other orthosteric ligands used in this study. The emission spectrum in the presence of HMA showed a higher relative intensity of the blue-shifted shoulder peak at 460 nm than the spectra recorded in the presence of either agonists or antagonists. The integrated fluorescence intensity measured in the presence of HMA was statistically indistinguishable from that observed for the agonist-bound A_2A_AR. These results imply that the allosteric ligand HMA stabilizes a distinct conformation of A_2A_AR, which is different from the conformations of both inactive apo or antagonist-bound A_2A_AR and active agonist-bound A_2A_AR. Thus, the employment of the A_2A_AR_L225C_-DyeC fluorescent sensor can report on conformational changes induced by A_2A_AR ligands with different modes of action.

### MD simulations of A2AARL225C-DyeC reveal the behavior of the label during receptor activation

In order to rationalize fluorescence changes observed upon A_2A_AR activation, we conducted MD simulations of A_2A_AR_L225C_-DyeC embedded in a lipid membrane both in the active and inactive receptor states. Since obtaining adequate structural ensembles of labeled conformers in unbiased MD simulations still represents a serious computational challenge [53], which is additionally complicated by the presence of a lipid bilayer, we employed an efficient enhanced sampling technique, known as metadynamics [54,55]. This approach allowed us to estimate free energy surfaces of DyeC attached to A_2A_AR_L225C_ and predict its preferred spatial positions in the active and inactive state of the receptor.

The estimated free energy surfaces revealed a remarkable difference in the predominant orientation of DyeC between the active and inactive conformations of A_2A_AR (Figure 4A–B). In the active state, regions with low free energy values were observed near the protein-lipid interface, where DyeC appears plunged into lipid headgroups. In contrast, the low free energy regions in the inactive state were located at the receptor surface, where the label is located in the vicinity of intracellular tips of protein helices remaining largely in the aqueous environment.

**Figure 4.**
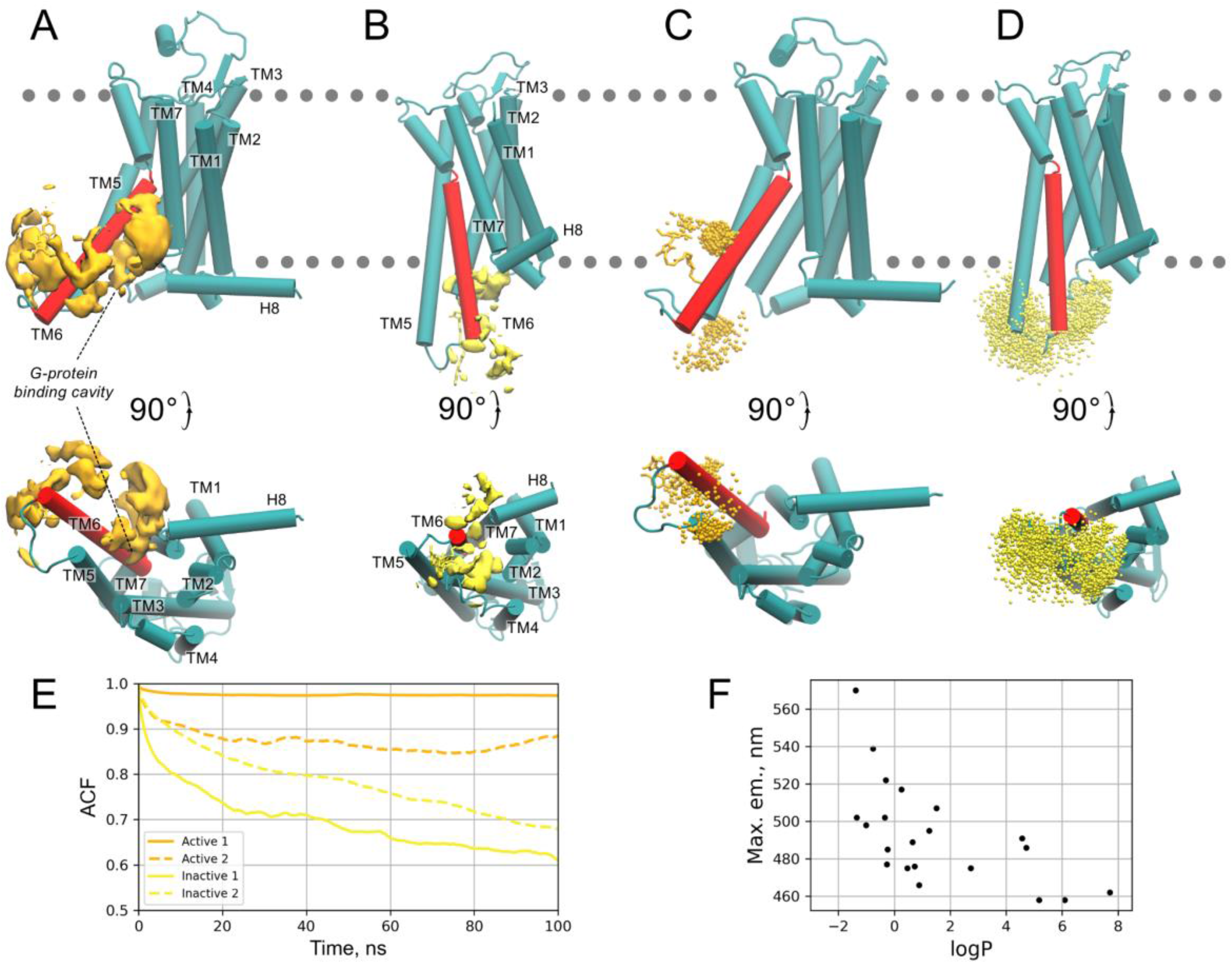
Molecular dynamics simulations of A2AARL225C-DyeC. **A-B:** The isosurfaces (yellow) delineate the low free energy regions (at +25 kJ/mol level relative to the global free energy minimum) explored by the Dye C label in the active (A) and inactive (B) states in metadynamics simulations. The A2AARL225C helices are labeled from TM1 to H8 (TM6 is colored in red), the G-protein binding site is labeled in the active state. **C-D:** Positions of the dimethoxybenzene ring of the DyeC label in the active (C) and inactive (D) complexes throughout the unbiased (i.e., without any external forces applied) MD simulations shown every 0.1 ns as orange/yellow dots, respectively. Each system was simulated for 1000 ns in two replicates. The positions of the lipid head groups are schematically indicated by the gray dotted line. **E:** Autocorrelation functions (ACF) calculated for a vector describing the DyeC label position in the unbiased simulations shown in panels C-D; **F:** Correlation between fluorescence emission maximum of DyeC in different solvents and their partition coefficient, logP. The logP values were obtained from the Pubchem/Chemeo databases [83,84].

We further conducted unbiased molecular simulations without applying any external forces to evaluate the mobility of the DyeC label in the uncovered free energy minima for both active and inactive states. The autocorrelation function (ACF) calculated for the vector describing the orientation of the label unveiled its significantly slower dynamics in the active state, in which DyeC was immersed in the lipid headgroup region or trapped between TM5 and TM6 helices of A_2A_AR, compared to the inactive state featuring the solvent-exposed DyeC conformation (see Figure 4C,D, and F).

These MD results allowed us to propose a mechanistic explanation for the changes in the spectral properties observed upon A_2A_AR activation. In the active state of the receptor, DyeC moves from an aqueous environment to the head group region of the lipid bilayer. This region has higher polarity due to the presence of charged phosphate and choline groups. The latter may be the reason for the red shift of the fluorescence emission maximum, which is consistent with the observed dependence of the DyeC emission maximum on the solvent logP (see Supplementary Table 1 and Figure 4G, compared with the consensus logP prediction = −2.65 obtained using the SwissADME service [59] for the zwitterionic phosphocholine molecule mimicking the POPC head group). In turn, in the active state, DyeC is more confined due to its encapsulation into the G-protein-binding cavity or interaction with TM6 (see Figure 4E). This is reflected in lower ACF and invokes a restriction of the intramolecular rotation (RIR) of DyeC that is known to be responsible for FQY enhancements in GFP-derived chromophores [60]. A concomitant fluorescence enhancement may be induced by the higher viscosity of the surrounding lipid environment (up to 5 orders of magnitude [56–58]) compared to the bulk water.

## Discussion

In this study we utilized solvatochromic dyes inspired by the GFP chromophore to label Rec and A_2A_AR and measured their responses to ligands. Two out of four dyes (DyeB and DyeC) exhibited Ca^2+^-dependent changes in their emission spectra when attached to the native C39 of Rec. Considering the structural similarity between Rec and other neuronal calcium sensors, DyeB and DyeC can be implemented for monitoring conformational changes of these proteins, including those involved in cardiac arrhythmia, Alzeheimer’s, Parkinson’s and Huntington’s diseases, and proliferation of cancer cells [61]. DyeC showed the most significant spectral change and was selected for A_2A_AR labeling. We specifically attached DyeC to the intracellular tip of TM6 of A_2A_AR containing a single point mutation (L225^6.27^C) using a crude membrane labeling procedure. We embedded the labeled A_2A_AR into ND to mimic a membrane-like environment. These measures improved the structural integrity of the receptor and minimized potential artifacts. Overall, our approach compared favorably with previously reported methods involving environmentally sensitive dyes (Supplementary table S3), where receptors were extensively mutated (up to six point mutations) or/and solubilized in detergents.

A_2A_AR_L225C_-DyeC exhibited a 20% increase in the integrated fluorescence intensity and a 5-nm red-shift in the fluorescence emission maximum in response to the interaction with full agonists (NECA and Adenosine) but showed no response to the interaction with antagonists (ZM241385 and SCH58261). Both the integrated fluorescence intensity and the I_520_/I_460_ ratio can be effectively used for quantitative description of the observed spectral changes. The latter ratiometric approach has the advantage that the readout does not depend on the receptor concentration. It can be also useful when analyzing the effects of allosteric ligands, such as HMA, which has a unique impact on the shape of the DyeC fluorescence spectra and can be distinguished from that caused by agonists. These spectral effects can be distinguished through ratiometric analysis but not by integrated fluorescence intensity. The distinctive shape of the emission spectrum implies that structural changes in A_2A_AR induced by HMA differ from those caused by orthosteric agonists. This finding aligns with a previous NMR study [32], which showed that HMA increases the population of a state distinct from those stabilized by NECA.

In terms of the magnitude of agonist-induced changes in the fluorescence intensity and the emission maximum wavelength, DyeC is competitive to other environmentally sensitive labels that have been previously employed to monitor GPCR activation (Supplementary Table S2). Most of the previous GPCR activation studies using fluorescent labels have been focused on the β_2_-adrenergic receptor (β_2_AR) that exhibits agonist-induced structural changes similar to A_2A_AR [62]. Dyes of various chemical natures were employed to label β_2_AR at positions homologous or close to the labeling position at the intracellular tip of TM6 used in our study. Among these dyes, bimane was the most common label that showed a 50% intensity change and a 15-nm spectral shift in the presence of agonists [63], which exceeded the fluorescence changes observed for DyeC in our experiments (20% intensity change and 5-nm spectral shift). However, bimane has an absorption spectrum with the maximum at a shorter wavelength (∼450 nm compared to 515 nm for DyeC) and a lower extinction coefficient (∼5,000 M^-1^cm^-1^ [64], compared to 14,000 M^-1^cm^-1^ for DyeC). Thus, despite the superior response characteristics of bimane, the red-shifted spectrum and the higher extinction coefficient of DyeC make it a more favorable dye for GPCR labeling in some applications.

Our MD simulations revealed that the observed changes in the DyeC fluorescence spectrum can be explained by the relocation of the fluorescent label to a more confined conformational space with a more hydrophilic environment upon receptor activation. This effect is caused by the agonist-induced outward movement of the DyeC-labeled intracellular tip of TM6, representing a common feature of the activation mechanism in class A GPCRs [65]. Considering these observations, we propose that our approach could be applied to other GPCRs enabling future research on their conformational dynamics.

## Methods

### Synthesis of dyes

Commercially available reagents of highest purity were used without additional purification. Merck Kieselgel 60 was used for flash chromatography. Thin layer chromatography was performed on silica gel 60 F_254_ glass-backed plates (Merck). Visualization was enabled by staining with KMnO_4_ and illumination with UV light (254 or 312 nm).

NMR spectra were recorded on a 700 MHz Bruker Avance NMR at 303 K and a 800 MHz Bruker Avance III NMR at 333 K. Chemical shifts are reported relative to the residue peaks of CDCl_3_ (7.27 ppm for ^1^H and 77.0 ppm for ^13^C) or DMSO-d_6_ (2.51 ppm for ^1^H and 39.5 ppm for ^13^C). Melting points were measured on a SMP 30 apparatus (Stuart Scientific). High-resolution mass spectra (HRMS) were recorded on a Bruker micrOTOF II instrument using electrospray ionization (ESI). The measurements were carried out in a positive ion mode (interface capillary voltage – 4500 V) or in a negative ion mode (3200 V); mass range from m/z 50 to m/z 3000; external or internal calibration was done with ESI Tuning Mix (Agilent). A syringe injection was used for solutions in acetonitrile, methanol, or water (flow rate 3 mL/min). Nitrogen was applied as a dry gas; interface temperature was set at 180 °C.

Detailed synthetic procedures in Supplementary Methods.

### Rec and RecC39D expression, purification and labeling

#### DNA constructs

The genetic construct for bacterial expression of Rec from Bos taurus (UniProt ID P21457) was prepared by inserting corresponding cDNA into a pET11d vector (Novagen, USA) between NcoI and BamHI restriction sites under the control of the T7 phage promoter as previously described [66]. The construct for the expression of the cysteine-free Rec mutant C39D (Rec_C39D_) was obtained by site-directed mutagenesis using the bovine Rec gene in a pET-11d plasmid as a template as previously described [67].

#### Rec and RecC39D expression

Rec and Rec_C39D_ were produced in *Escherichia coli* strain BL21-CodonPlus®(DE3)-RIL-X (Agilent, USA) co-transformed with a pBB131 plasmid, carrying a gene for the N-myristoyl transferase-1 from Saccharomyces cerevisiae (UniProt ID P14743). To obtain the myristoylated form of Rec, cells were cultivated for 5 h (250 rpm, 37 °C) in the presence of myristic acid (20 mg/L) added to the medium immediately prior to the induction of the expression with 0.5 mM isopropyl β-D-1-thiogalactopyranoside. Rec/Rec_C39D_-containing fractions were obtained by exposing cells to repetitive freeze-thaw cycles in lysis buffer (50 mM Tris-HCl, pH=7.5, 5 mM MgCl_2_, 1 mM EDTA, 0.1 mM PMSF, 3 mM DTT) and treatment of the resulting suspension with lysozyme (25–50 µg/mL, 20 min, on ice) with subsequent centrifugation (12,000 ×g, 20 min, 4 °C).

#### Rec and Rec_C39D_ purification

For the primary isolation of Rec and Rec_C39D_, the obtained fractions were mixed on ice with calcium chloride (3 mM), loaded onto a Phenyl Sepharose column (Cytiva, USA), equilibrated with a buffer (20 mM Tris-HCl, pH=7.5, 2 mM CaCl_2_, 2 mM MgCl_2_ and 1 mM DTT), and the target proteins were eluted with the same buffer containing 1 mM EGTA instead of CaCl_2_. For the final purification and concentration of recoverin forms, the EGTA fractions were loaded onto a HiTrap Q FF column (Cytiva, USA) equilibrated with a buffer (20 mM Tris-HCl, pH=7.5, 1 mM DTT), and the proteins were eluted using a linear salt gradient from 0 to 1 M NaCl. After the purification, Rec and Rec_C39D_ were dialyzed against a storage buffer (20 mM Tris-HCl, pH=7.5, 1 mM DTT), aliquoted, and stored at –80 °C (Supplementary Figure S6). The degree of Rec myristoylation in the obtained preparations was determined by analytical HPLC using Luna C18 reversed-phase column (Phenomenex) in the acetonitrile-water system and was more than 95% (Supplementary Figure S7). The final yield was 25-30 mg of the Rec/Rec_C39D_ per liter of bacterial culture.

#### Rec labeling

Rec samples were transferred to a labeling buffer (50 mM HEPES, pH=7.0, 5 mM EGTA) using 10 kDa MWCO Amicon Ultra centrifugal filter unit (Merck Millipore, USA), and concentrated to 0.5 μg/mL. Twenty-fold molar excess of each of the dyes (DyeA, DyeB, DyeC or DyeD) dissolved in DMSO was then added to Rec preparation, yielding the final DMSO concentration of 6% (v/v). The mixture was incubated for 16 h at 4 °C with constant rotation on an orbital shaker (15 rpm) in the dark. The unbound fluorescent labels were washed out with a Rec-washing buffer (50 mM HEPES pH=8.0) in a Stirred Cell Model 8003 (Millipore), with a 10 kDa MWCO regenerated cellulose filter (Millipore). The protein concentration and labeling efficiency were calculated from the absorption spectra as described below (see “Labeled proteins absorption and emission measurements”).

### A2AAR expression, purification, and labeling

#### DNA construct

The nucleotide sequence of the human *ADORA2A* gene encoding A_2A_AR (2-316 aa) (UniProt ID C9JQD8) was obtained from the cDNA Resource Center (cdna.org, #ADRA2A0000) and modified for expression in *Spodoptera frugiperda*. The final gene construct was complemented from the 5’-terminus by nucleotide sequences of the hemagglutinin signal peptide (MKTIIALSYIFCLVFA), FLAG-tag (DYKDDDDK), linker (AMGQPVGAP), and from the 3’-terminus by the 10×His-tag nucleotide sequence and inserted into a pFastBac1 vector (Invitrogen) via the BamHI(5’) and HindIII(3’) restriction sites. The L225^6.27^C mutation was introduced by PCR. A snake-plot representation of the protein construct is shown in Supplementary Figure S8.

#### A2AAR expression

A high-titer recombinant baculovirus (>10^9^ viral particles per mL) for A_2A_AR expression in insect cells (Sf9) was obtained following a modified Bac-to-Bac system protocol (Invitrogen). Recombinant baculoviruses were generated by transfecting 2.5 μg of a transfer bacmid into Sf9 cells (2.5 mL at a density of 10^6^ cells/mL) using 3 μL of X-tremeGENE™ HP DNA Transfection Reagent (Roche) and 100 μL Transfection Medium (Expression Systems). The cell suspension was incubated for 4 days with shaking using a Shel Lab incubator at 27 °C and 300 rpm in 24-deep well U-bottom plates covered with Breathe-Easy membranes (Greiner BioOne). A P0 viral stock was isolated by centrifugation at 2000 ×g for 5 min, and used to produce a high-titer baculovirus stock (P1): 40 mL of cells at 2 ×10^6^ cells/mL density were infected with 2.5 mL supernatant, and grew for 72 h with shaking at 27 °C and 120 rpm (Innova 44, New Brunswick). Sf9 cells at a cell density of 2–3 ×10^6^ cells/mL were infected with the P1 virus at the multiplicity of infection equal to 5. Expression was performed with shaking at 27 °C and 120 rpm (Innova 44, New Brunswick), in 1 L vent-cap flasks (Corning). Cells were harvested by centrifugation at 2,000 ×g for 10 min, after 48 h post infection, and stored at –80 °C until further use. Cell counts, viral titers and expression levels were measured by flow cytometry on a BD Accuri C6 (BD Biosciences).

#### A2AAR purification and labeling

The biomass obtained from 1 L of cell culture was thawed on ice by adding 200 µL of a protease inhibitor cocktail (500 µM AEBSF (Gold Biotechnology), 1 µM E-64 (Cayman Chemical), 1 µM leupeptin (Cayman Chemical), 150 nM aprotinin (AG Scientific)) in a low-salt buffer (10 mM HEPES pH=7.5, 10 mM MgCl_2_, 20 mM KCl) to 100 mL (scaled if necessary). The mixture was homogenized in a high-tight 100 mL Potter douncer on ice, centrifuged for 20 min at 4 °C and 220,000 ×g. The supernatant was discarded, and the pellet was resuspended in 100 mL of a high-salt buffer (10 mM HEPES pH=7.5, 10 mM MgCl_2_, 20 mM KCl, 1 M NaCl) supplemented with 100 µL protease inhibitor cocktail, centrifuged for 30 min at 4 °C and 220,000 ×g. The resuspension and centrifugation were repeated one more time.

Fluorescent labeling and all further procedures were carried out at 4 °C in the dark or under a dim red light and with constant rotation on an orbital shaker (15 rpm) during any incubation.

The washed membranes obtained from 1 L of cell culture were resuspended in 20 mL of a labeling buffer (20 mM HEPES pH=7.0, 10 mM MgCl_2_, 20 mM KCl, 4 mM theophylline, 50 µL protease inhibitor cocktail), mixed with a dye solution (4 mg of dye B or C, dissolved in 60 µL of DMSO), and incubated for 16 h. After labeling, membrane fractions were pelleted by ultracentrifugation at 220,000 ×g for 30 min and washed three times with the high-salt buffer to remove unbound fluorescent labels.

The labeled membranes were homogenized in 50 mL of solubilization buffer (50 mM HEPES pH=7.5, 800 mM NaCl, 5% v/v glycerol, 0.5/0.1% w/v DDM/CHS (Sigma), 4 mM theophylline (Sigma), 50 µL protease inhibitor cocktail). Receptor solubilization was carried out for 4 h. The insoluble debris was eliminated then by centrifugation for 1 h, 650,000 ×g, while the target protein remained in the supernatant. The supernatant was incubated with 500 µL of a TALON resin (Clontech) for 16 h.

The resin was washed with 10 column volumes (CV) of wash buffer 1 (50 mM HEPES pH=7.5, 800 mM NaCl, 10% v/v glycerol, 25 mM imidazole, 0.1/0.02% w/v DDM/CHS, 10 mM MgCl_2_, 8 g/mol ATP (Sigma), 4 mM theophylline), then 5 CV of wash buffer 2 (50 mM HEPES pH=7.5, 800 mM NaCl, 10% v/v glycerol, 50 mM imidazole, 0.05/0.01% w/v DDM/CHC, and 4 mM theophylline). The receptor was eluted with 3 CV of elution buffer (25 mM HEPES pH=7.5, 800 mM NaCl, 10% v/v glycerol, 220 mM imidazole, 0.01/0.002% w/v DDM/CHS, and 4 mM theophylline). The eluted receptor was desalted from imidazole using a size-exclusion PD10 column (GE Healthcare) equilibrated with the desalt buffer (25 mM HEPES pH=7.5, 800 mM NaCl, 0.01/0.002% w/v DDM/CHS).

The labeled and purified receptor was concentrated using a 30 kDa MWCO filter (Merck, Amicon Ultra) to 10–15 µM. Receptor purity and homogeneity were assessed by SDS-PAGE and analytical size-exclusion chromatography (SEC) accordingly (Supplementary Figures S9 and S10).

#### A2AAR reconstitution into ND

The Membrane Scaffold Protein 1D1 (MSP1D1) for ND was expressed in the *Escherichia coli* strain BL21(DE3) using a gene (nucleotide sequence was taken from McLean M.A. at al. [68] and synthesized de novo, Genescript) with an N-terminal 6×His-tag and a TEV-protease site cloned into a pET28a vector between the NcoI and HindIII restriction sites (Supplementary Table S3). MSP1D1 was purified using IMAC Ni-NTA Agarose (Qiagen) with further cleavage of 6×His-tag by TEV protease. The lipid mixture of POPC:POPG (Avanti Polar Lipids) in chloroform was prepared at a molar ratio 7:3. The lipid film was dried under a gentle nitrogen gas stream, followed by removal of the solvent traces under vacuum first with the use of a rotary evaporator (Hei-VAP Ultimate ML/G1, Heidolph) and then with deep vacuum oil-free pump (ChemStar® Dry Oil-Free Deep Vacuum System) overnight, next day solubilized in 100 mM sodium cholate (Sigma) and stored at –80 °C until further use.

The purified A_2A_AR (WT or L225С^6.27^) in DDM/CHS micelles was mixed with MSP1D1 and the POPG:POPC lipid mixture at a molar ratio A_2A_AR:MSP1D1:lipids=1:5:50. The final sodium cholate concentration was maintained in the range of 25–30 mM, the typical final receptor concentration was 0.5⎼0.6 mg/mL. After 1 h pre-incubation, the mixture was incubated overnight with wet Bio-Beads SM-2 (0.14 g of beads for 1 g of detergent were washed in methanol and equilibrated with 25 mM HEPES, pH=7.5, 150 mM NaCl). The next morning, a fresh portion of Bio-Beads for an additional 4 h incubation was added, beads were discarded and the supernatant containing reconstituted into ND A_2A_AR was incubated for 4 h with 250 µL of Ni-NTA resin (Qiagen) for separating from empty ND. The protein was eluted in the elution buffer (25 mM HEPES pH=7.5, 150 mM NaCl, 150 mM imidazole), and then desalted from imidazole using size-exclusion PD-10 column equilibrated with desalt ND buffer (25 mM HEPES pH=7.5, 150 mM NaCl).

A_2A_AR reconstituted into ND was concentrated using a 100 kDa MWCO filter (Merck, Amicon Ultra) to 10-15 µM. Labeling efficiency was calculated from the absorption spectrum measured as described below (see “Labeled proteins absorption and emission measurements”).

#### SDS-PAGE and SEC

The samples were subjected to SDS-PAGE using a Mini-Protean IV system (Bio-Rad) with polyacrylamide gel (5% concentrating gel with AA:bisAA ratio of 29:1 and 15% resolving gel with 19:1 AA:bisAA ratio). The protein was loaded in the amount of 5 μg of receptor per lane previously mixed with a loading buffer (25 mM Tris pH=6.8, 25% v/v glycerol, 0.25% SDS, bromophenol blue) without preheating and stained after separation with Coomassie Brilliant Blue R-250.

Analytical size-exclusion chromatography was performed on a Dionex Ultimate 3000 instrument (Thermofisher) equipped with a Nanofilm Sec 250 (Sepax technologies, cat# 201250-4625) gel filtration analytical column. The column was equilibrated with chromatographic buffer: 0.05/0.01% w/v DDM/CHC, 25 mM HEPES pH=7.5, 500 mM NaCl, 20 mM MgCl_2_, 2% v/v glycerol. For ND, no detergents were added in the chromatographic buffer. The flow rate was 0.35 mL/min, protein absorption was detected at 280 nm, and 40 µL of the sample was injected.

### Free dye absorption and emission measurements

UV-VIS spectra of free dyes were recorded on a Varian Cary 100 (Agilent Varian) spectrophotometer. Fluorescence excitation and emission spectra were recorded on a Cary Eclipse (Agilent Technologies) fluorescence spectrophotometer.

The fluorescence quantum yields were calculated according to the procedure described in the literature [69] with the use of Coumarine 153 as a standard. The quantum yield was calculated by the formula:

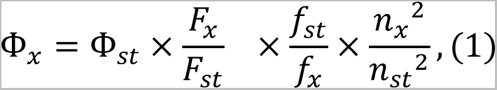

where *F* is the area under the emission peak, *f* is the absorption factor (see below), *n* is the refractive index of the solvent, *Φ* is the fluorescence quantum yield, the subscript *x* corresponds to the dye of interest, the subscript *st* – for standards.

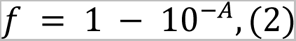

where *A* is the absorbance at the excitation wavelength.

The molar extinction coefficient was calculated by the formula:

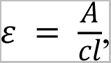

where *A* is the absorbance maxima, *c* is the molar concentration, *l* is the pathlength.

### Absorption and emission measurements of labeled proteins

All absorption and emission measurements for both labeled A_2A_AR and Rec were carried out at 25 °C on a Synergy H4 (Agilent BioTek) plate reader with Take3 Microvolume Plates; 2.5 µL of sample was added in each microspot. After measurements, the spectrum of the buffer was measured in each microspot and subtracted from the sample spectra. All obtained data were analyzed using the GraphPad Prizm 9.4.1 software.

#### Concentration and labeling efficiency calculations

Protein concentrations were calculated from absorption at 280 nm using the molar extinction coefficients of 24,175 M^-1^cm^-1^ (Rec), 51,880 M^-1^cm^-1^ (A_2A_AR), 18,200 M^-1^cm^-1^ (MSP1D1) and the light pathlength of 0.05 cm.

Labeling efficiencies for proteins with DyeA, DyeB, DyeC, and DyeD were estimated by measuring their absorbance at 420, 440, 390, and 390 nm using the extinction coefficients of 19,000 M^-1^cm^-1^, 21,000 M^-1^cm^-1^, 14,000 M^-1^cm^-1^ and 15,000 M^-1^cm^-1^, respectively [3].

#### Labeled Rec emission measurements

1 mM CaCl_2_ and 1 mM EGTA were added separately to the labeled protein in Rec-washing buffer (50 mM HEPES pH=8.0) with final concentrations of 100 μM for CaCl_2_ or EGTA in a total volume of 9 µL: Rec concentration was equal between two protein samples labeled with same dye and varied in a range of 2-10 μM for samples labeled with different dyes. After 20 min of incubation at 4 °C in the dark, fluorescence emission spectra of Rec with DyeA, DyeB, and DyeCwere then measured with excitation at 440 nm, 460 nm, and 410 nm, respectively, and a 9 nm emission bandwidth.

#### Labeled A2AAR emission measurements

Ligands (ZM241385 (Cayman Chemical), SCH58261 (Tocris), Adenosine (Tocris), HMA (Merck) or NECA (Tocris)) were dissolved in DMSO to make 100 mM stock solutions. Then 1 mM and 100 μM stock solutions of ligands in the desalt ND buffer were prepared. The samples with labeled A_2A_AR (WT or L225^6.27^C) and 100 μM of a ligand (or no ligand for apo sample) were prepared in the desalt ND buffer in a final volume of 9 µL. The final protein concentrations were equal between different ligands within the same run of the measurements, but varied in a range of 2-10 μM for different receptor batches.

After 20 min incubation at 4 °C in the dark, emission spectra of A_2A_AR with DyeB and DyeC were measured with excitation at 460 nm and 410 nm, respectively, and a 9 nm emission bandwidth.

Emission spectra for each protein sample from at least three different protein purifications were recorded with at least three technical repeats for each purification. To compensate for differences of protein concentration between different purification batches, all measured fluorescence spectra were normalized to the maximum of the fluorescence spectra of A_2A_AR_L225C_-DyeC with NECA measured in the same run and averaged over at least three technical replicas.

#### Displacement experiments

Two experiments were conducted to demonstrate the reversibility of conformational changes upon ligand displacement. In the beginning of each experiment, three samples of apo A_2A_AR_L225C_-DyeC in the desalt ND buffer were prepared, 1 µg of receptor in each. The initial volumes of 8.1 µL, 7.8 µL and 6.9 µL were selected to amount to equal final volumes of 9 µL after all further additives. The first experiment was carried out to replace NECA with ZM241385. 0.9 µL of 100 µM NECA were added to each of the three A_2A_AR_L225C_-DyeC samples. After 20 min incubation at 4 °C, an emission spectrum using excitation at 410 nm was measured for the first sample containing 10 µM of NECA. Then 0.27 µL of 1 mM ZM241385 were added to the remaining two samples. After 20 min incubation at 4 °C, the emission spectrum of the second sample containing 10 µM of NECA and 30 µM of ZM241385 was measured. Finally 0.9 µL of 1 mM NECA was added to the remaining third sample to final cumulative concentration of 30 µM of ZM241385 and 110 µM of NECA. The emission spectrum of the third sample was measured after 20 min incubation at 4 °C. In the second experiment, the same steps were repeated, but the order of adding ZM241385 and NECA was opposite. The final protein concentration was 2 µM in each measured sample.

### Molecular dynamics simulations

Protein models were prepared using the structure of a thermostabilized A_2A_AR in complex with ZM241385 (PDB ID: 3PWH, [70]) and the structure of A_2A_AR in complex with mini-Gs (PDB ID: 5G53, [71]) for inactive and active states, respectively. The thermostabilized mutations were mutated back to the native amino acids and the missing regions were added using MODELLER [72]. All ionizable amino acids were modeled in their standard ionization state at pH=7.

The DyeC label was attached to the position L225^6.27^C by aligning its backbone moiety with the corresponding group of the original amino acid mutated to cysteine. Both active and inactive protein models were embedded in a lipid bilayer consisting of POPC and solvated using the CHARMM-GUI server [73]. The final models contained 203 POPC lipids, 21,674 water molecules, 86/95 Na^+^/Cl^−^ ions in the active state (97,334 atoms in total, box dimensions 9.09 × 9.09 × 11.48 nm), and 205 POPC lipids, 25,406 water molecules, 98/107 Na^+^/Cl^−^ ions in the inactive state (108,682 atoms in total, box dimensions 8.73 × 8.73 × 14.04 nm).

For each state, we produced two simulation replicates starting from two alternative orientations of DyeC (Supplementary Figure S5) to overcome the problem of slow DyeC flipping from one side of the TM5-ICL3-TM6 fragment to the other side. In order to generate the two alternative starting conformations with the fluorescent label oriented on the opposite sides of the TM5-ICL3-TM6 fragment (i.e., pointing towards and outwards the protein center), we run short steered MD simulations with the moving harmonic potential (force constant of 1000 kJ/mol/nm^2^) applied to one of the carbon atoms of the dimethoxybenzene ring pulling it away from the protein center-of-mass with the constant velocity of 10^-4^ nm/ps. These steered simulations were conducted until the label flipped across ICL3 (∼30 ns) and resulted in four starting conformations, i.e., two alternative label conformations for each state, active and inactive. The starting models were subjected to the standard CHARMM-GUI equilibration comprising a steep descent minimization followed by several short equilibration simulations, ∼2 ns in total, with the harmonic restraints applied to the protein and lipids.

After equilibrating the starting models according to the protocol described above, we performed metadynamics MD simulations to estimate the three-dimensional free energy surfaces for the fluorescent label in the active/inactive states. Metadynamics is an enhanced sampling technique, which allows to reconstruct free energy profiles/surfaces along selected collective variable(s) by adding numerous repulsive potentials or “hills” (typically, Gaussian-shaped) to the total potential of the system forcing the latter to explore its configurational space faster and broader [54,55]. We have chosen three projections of the vector connecting the Cα atom of the label with the proximal carbon atom of its dimethoxybenzene ring onto the three Cartesian axes as the collective variables biased in the metadynamics simulations since they provide a straightforward and efficient way to describe the general orientation of the fluorescent label. Gaussian hills with the width of 0.1 nm and the height of 0.25 kJ/mol were added every 0.5 ps. The grid was used to efficiently store the accumulated hills. Each metadynamics simulation was run for 500 ns. The convergence of the simulations was estimated by calculating the volume explored by the label using the custom script available at https://github.com/porekhov/A2a_GFP_core_dyes. The cumulative explored volume plotted as a function of simulation time is shown in Supplementary Figure S5.

Throughout the equilibration and metadynamics simulations, positional restraints were applied to Cα atoms of the transmembrane region of A_2A_AR as well as to the intracellular segment of TM6 encompassing the labeling position 225 and its neighboring amino acids to preserve the protein state and to prevent fluctuations of the label’s Cα atom, which was used as the reference point for calculation of the collective variables. Apart from it, we also applied dihedral restraints (force constant of 1000 kJ/mol/rad^2^) to the backbone atoms of the solvent-exposed protein regions in the α-helical conformation.

Since the relocation of the fluorescent label from one side of the TM5-ICL3-TM6 fragment to the opposite side is a relative slow process particularly hindered by the restraints applied to the protein backbone and limiting the thorough sampling of label orientations, we run duplicate metadynamics simulations for each state starting from two alternative conformations of the label obtained as noted above.

Additionally, four unbiased simulations (i.e., without any external forces applied to a system) were run to estimate mobility of the DyeC label in the active and inactive A_2A_AR states starting from two alternative label orientations similar to the metadynamics simulations (see above). Each simulation was carried out for 1000 ns. The autocorrelation function was calculated as a measure for the mobility of fluorescent label using the rotacf tool in GROMACS according to the following expression, ACF(*t*) = <**v**(τ)⋅**v**(τ+*t*)>_τ_, where **v** is a vector describing the orientation of the fluorescent label and defined by Cα and the proximal carbon atom of the dimethoxybenzene ring of DyeC.

All MD simulations were performed by GROMACS version 2022.3 [74] with the PLUMED plugin, version 2.8.1 [75] allowing for metadynamics. A time step of 1-2 fs was used for equilibration simulations (3 × 125 ps with the 1-fs time step followed by 3 × 500 ps with the 2-fs time step), while all production metadynamics and unbiased simulations were performed with a 5 fs time step allowed by repartitioning the mass of heavy atoms into the bonded hydrogen atoms [76] and the LINCS constraint algorithm [77]. Production simulations were run in NVT ensemble with v-rescale thermostat (τ_T_ = 1.0 ps, T_ref_ = 303.15 K), Van-der-Waals interactions were treated using the cut-off scheme, electrostatics – using PME.

The Amber99SB-ILDN force field was used for the protein, lipids, and ions [78] along with the TIP3P water model. The topology for the fluorescent label was obtained using Antechamber [79] with some dihedral parameters adopted from [80]. Conformation of the label was preliminary optimized at the DFT level with B3LYP functional and Def2-SVP/Def2/J basis set in Orca [81]. The resulting topology and structure files are available in the Supplementary Materials.

## Acknowledgements

This work is supported by RSCF research grant 22-74-00024. J.Ho. acknowledges support from the Flemish government through long-term structural funding Methusalem (CASAS2, Meth/15/04).

## Author contributions statement

N.B., V.M. and S.K. synthesized the dyes and measured their optical and NMR spectra under supervision of Alr.M., M.B. and K.M.

D.Z. expressed and purified Rec under supervision of E.Z.

A.Be. and P.Kh. expressed, purified, and reconstituted A_2A_AR in NDs under supervision of P.Ku. and Alx.M.

A.Be., I.M. and A.Bo. labeled Rec and A_2A_AR, measured and analyzed absorption and fluorescence spectra of the labeled proteins under supervision of T.G. and V.B.

P.O. performed and analyzed MD simulations

A.Be., I.M., P.O. and V.B. prepared the draft of the manuscript A.Be., M.B., Alr.M., I.M. and V.B. conceived the study

A.Be., I.M., P.O., E.Z., V.I., S.P., J.Ho, J.He, V.C., T.G., M.B. and V.B. discussed the data, analysis, and contributed to writing the manuscript

V.B. supervised the work

All the authors contributed to analyzing the data and editing the manuscript

