## Supplemental Information for "Monitoring GPCR Conformation with GFP-Inspired Dyes"

### Table of content

**Synthetic procedures	2**

**Supplemental Figures	10**

Supplementary Figure S1. Absorbance spectra of labeled Rec with all used dyes.	10

Supplementary Figure S2. Absorbance spectra of labeled A_2A_AR.	11

Supplementary Figure S3. Fluorescence emission spectra of A_2A_AR_WT_-DyeC with added ligands.	12

Supplementary Figure S4. Emission spectra of A_2A_AR_L225C_-DyeC after sequential addings of NECA and ZM241385.	13

Supplementary Figure S5. Metadynamics convergence analysis.	14

Supplementary Figure S6. SDS-PAGE (12.5%) of fractions obtained during purification of Rec/Rec_C39D_.	15

Supplementary Figure S7. Analytical HPLC of Rec and Rec_C39D_ in the acetonitrile-water system.	16

Supplementary Figure S8. Snake-plot presentation of protein construct of A_2A_AR.	17

Supplementary Figure S9. SDS-PAGE of A_2A_AR-DyeC samples.	18

Supplementary Figure S10. SEC of A_2A_AR in micelles and nanodiscs.	19

Supplementary Figure S11. Structures of DyeB maleimide isomers.	20

Supplementary Figure S12. Structures of DyeC maleimide isomers.	21

Supplementary Table S1. Solvatochromic properties of maleimide compounds DyeA, DyeB, DyeC, and DyeD.	22

Supplementary Table S2. I520/I460 ratios for A_2A_AR_L225C_-DyeC with various ligands.	23

Supplementary Table S3. Detection of GPCR structural changes with environmentally sensitive dyes.	24

**References	25**

### Synthetic Procedures

**Preparation of (Z)-4-(4-benzylidene-2-methyl-5-oxo-4,5-dihydro-1H-imidazol-1-yl)butanoic acids**

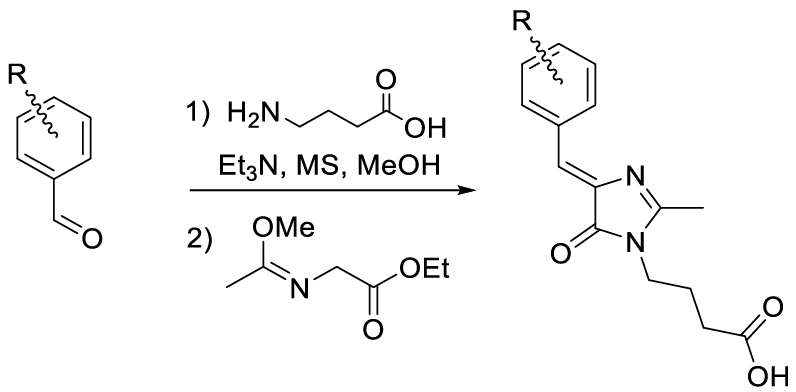

The corresponding aromatic aldehyde (12 mmol) was dissolved in MeOH (50 mL) and mixed with 4-aminobutanoic acid (1.35 g, 13 mmol), triethylamine (2.8 mL, 20 mmol) and MS 4Å (5 g) and 3Å (5 g). The mixture was stirred for 5 days at room temperature. The mixture was filtered; MS were washed with MeOH (2 × 10 mL). The solvent was evaporated and ethyl((methoxy)amino)acetate (2.4 g, 15 mmol) was added to the residue. The mixture was stirred for 4 days at room temperature, solvents were evaporated and the product was purified by column chromatography (CH_2_Cl_2_-iPrOH-AcOH, 100/5/0.5).

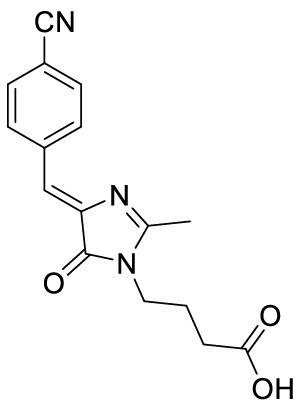

**(*Z*)-4-(4-(4-cyanobenzylidene)-2-methyl-5-oxo-4,5-dihydro-1*H*-imidazol-1-yl)butanoic acid**

Yellow solid (1.92 g, 54%); mp = 176-179 ºС; ^1^H NMR (700 MHz, DMSO-d_6_) δ ppm 12.10 (br. s., 1 H), 8.38 (d, J=8.4 Hz, 2 H), 7.90 (d, J=8.4 Hz, 2 H), 7.02 (s, 1 H), 3.61 (t, J=7.2 Hz, 2 H), 2.41 (s, 3 H), 2.27 (t, J=7.2 Hz, 2 H), 1.79 (quin, J=7.2 Hz, 2 H); ^13^C NMR (176 MHz, DMSO-d_6_) δ ppm 173.8, 169.8, 166.2, 140.9, 138.6, 132.3, 132.0, 122.0, 118.7, 111.3, 39.5, 30.7, 23.8, 15.4; HRMS (ESI) m/z: 298.1189 found (calcd for C_16_H_16_N_3_O_3_^+^, [M+H]^+^ 298.1186).

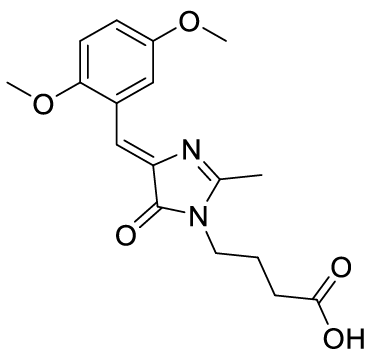

**(*Z*)-4-(4-(2,5-dimethoxybenzylidene)-2-methyl-5-oxo-4,5-dihydro-1*H*-imidazol-1-yl)butanoic acid (DyeC acid)**

Yellow solid (2.51 g, 63%); mp = 148-151 ºС with decomposition; ^1^H NMR (700 MHz, DMSO-d_6_) δ ppm 12.10 (br. s., 1 H), 8.39 (s, 1 H), 7.26 (s, 1 H), 7.23-7.28 (m, 2 H), 3.83 (s, 3 H), 3.74 (s, 3 H), 3.59 (t, J=7.2 Hz, 2 H), 2.38 (s, 3 H), 2.26 (t, J=7.2 Hz, 2 H), 1.78 (quin, J=7.2 Hz, 2 H); ^13^C NMR (176 MHz, DMSO-d_6_) δ ppm 173.8, 170.0, 163.8, 153.0, 152.9, 138.3, 122.9, 117.6, 117.1, 117.0, 112.2, 56.1, 55.4, 39.3, 30.7, 23.9, 15.4; HRMS (ESI) m/z: 333.1452 found (calcd for C_17_H_21_N_2_O_5_^+^, [M+H]^+^ 333.1445).

***Preparation of 4-(4-((Z)-benzylidene)-2-((E)-styryl)-5-oxo-4,5-dihydro-1H-imidazol-1-yl)butanoic acids***

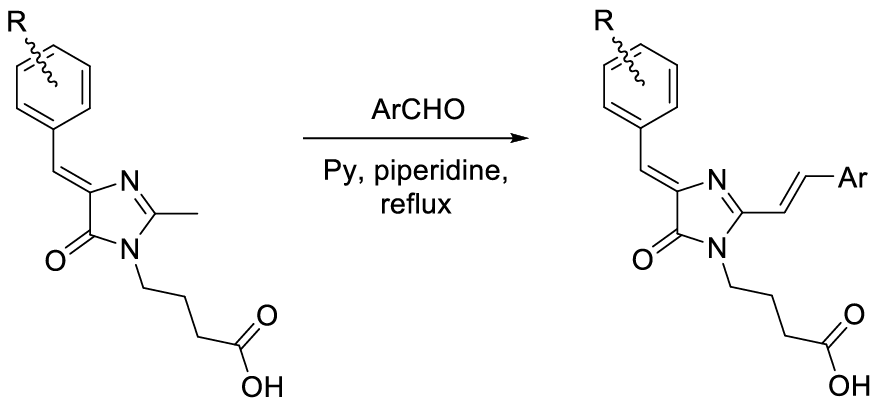

To the solution of (Z)-4-(4-benzylidene-2-methyl-5-oxo-4,5-dihydro-1H-imidazol-1-yl)butanoic acid (1 mmol) in pyridine (5 mL) piperidine (0.01 mL) and corresponding aldehyde (5 mmol) were added. The mixture was refluxed for 24 h and the solvent was evaporated. The residue was dissolved with mixture EtOAc (200 mL) and AcOH (1 mL), washed brain (2 × 10 mL) and dried over Na_2_SO_4_. The solvent was evaporated and the product was purified by column chromatography (CH_2_Cl_2_-MeOH, 100/5).

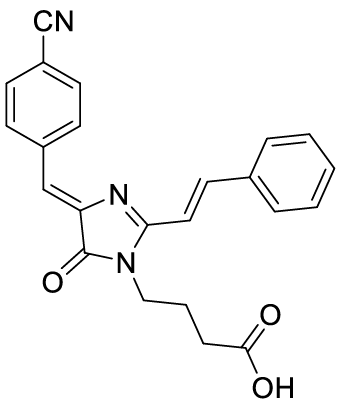

**4-(4-((*Z*)-4-cyanobenzylidene)-2-((*E*)-styryl)-5-oxo-4,5-dihydro-1*H*-imidazol-1-yl)butanoic acid (DyeA acid)**

Yellow solid (275 mg, 71%); mp = 194-197 ºС; ^1^H NMR (700 MHz, DMSO-d_6_) δ ppm 12.29 (br. s., 1 H), 8.48 (d, J=8.2 Hz, 2 H), 8.15 (d, J=15.6 Hz, 1 H), 7.87 - 7.93 (m, 4 H), 7.45 - 7.52 (m, 3 H), 7.33 (d, J=15.6 Hz, 1 H), 7.08 (s, 1 H), 3.81 (t, J=7.3 Hz, 2 H), 2.32 (t, J=7.0 Hz, 2 H), 1.80 (quin, J=7.2 Hz, 2 H); ^13^C NMR (176 MHz, DMSO-d_6_) δ ppm 174.1, 169.8, 162.1, 142.0, 141.5, 139.0, 134.9, 132.3, 132.3, 130.5, 129.0, 128.6, 122.0, 118.8, 113.4, 111.2, 38.9, 30.6, 24.5; HRMS (ESI) m/z: 386.1510 found (calcd for C_23_H_20_N_3_O_3_^+^, [M+H]^+^ 386.1499).

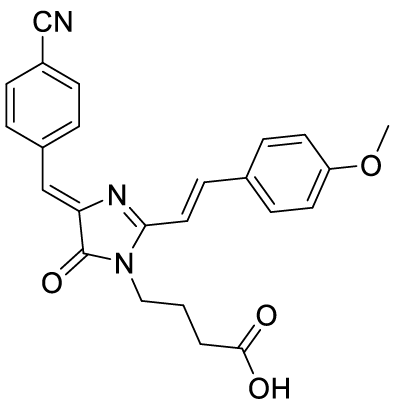

**4-(4-((*Z*)-4-cyanobenzylidene)-2-((*E*)-4-methoxystyryl)-5-oxo-4,5-dihydro-1*H*-imidazol-1-yl)butanoic acid (DyeB acid)**

Orange solid (310 mg, 81%); mp = 217-220 ºС; ^1^H NMR (700 MHz, DMSO-d_6_) δ ppm 8.46 (d, J=8.2 Hz, 2 H) 8.11 (d, J=15.6 Hz, 1 H), 7.86-7.90 (m, 4 H), 7.20 (d, J=15.6 Hz, 1 H), 7.04 (d, J=8.8 Hz, 2 H), 7.01 (s, 1 H), 3.84 (s, 3 H), 3.79 (t, J=7.3 Hz, 2 H), 2.27 (t, J=7.0 Hz, 2 H), 1.78 (quin, J=7.2 Hz, 2 H); ^13^C NMR (176 MHz, DMSO-d_6_) δ ppm 169.9, 162.5, 161.4, 142.0, 141.7, 139.2, 132.3, 132.1, 130.6, 127.6, 120.9, 118.9, 114.5, 114.5, 110.9, 110.6, 55.4, 39.0, 31.3, 24.8; HRMS (ESI) m/z: 416.1609 found (calcd for C_24_H_22_N_3_O_4_^+^, [M+H]^+^ 416.1605).

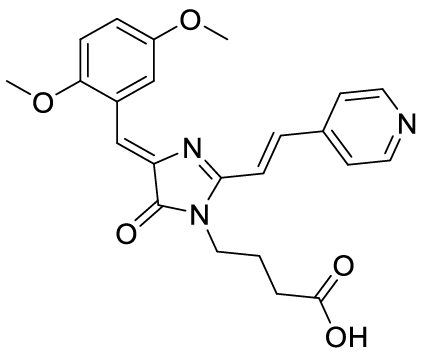

**4-(4-((*Z*)-2,5-dimethoxybenzylidene)-2-((*E*)-2-(pyridin-4-yl)vinyl)-5-oxo-4,5-dihydro-1*H*-imidazol-1-yl)butanoic acid (DyeD acid)**

Orange solid (290 mg, 69%); mp = 254-257ºС; ^1^H NMR (700 MHz, DMSO-d_6_) δ 12.21 (br. s., 1 H), 8.67 (d, J=5.9 Hz, 2 H), 8.54 (s, 1 H), 7.93 (d, J=15.8 Hz, 1 H), 7.79 (d, J=5.9 Hz, 2 H), 7.54 (d, J=15.6 Hz, 1 H), 7.42 (s, 1 H), 7.03-7.08 (m, 2 H), 3.86 (s, 3 H), 3.79 - 3.84 (m, 5 H), 2.31 (t, J=7.1 Hz, 2 H), 1.80 (quin, J=7.2 Hz, 2 H); ^13^C NMR (201 MHz, DMSO-d_6_) δ ppm 174.0, 169.8, 159.2, 153.4, 153.0, 150.3, 141.9, 138.7, 137.5, 123.1, 122.0, 119.2, 118.4, 118.3, 116.3, 112.4, 56.1, 55.3, 38.8, 30.4, 24.5; HRMS (ESI) m/z: 422.1719 found (calcd for C_23_H_24_N_3_O_5_^+^, [M+H]^+^ 422.1710).

***Preparation of 4-(4-((Z)-benzylidene)-5-oxo-4,5-dihydro-1H-imidazol-1-yl)-N-(2-(2,5-dioxo-2,5-dihydro-1H-pyrrol-1-yl)butanamide***

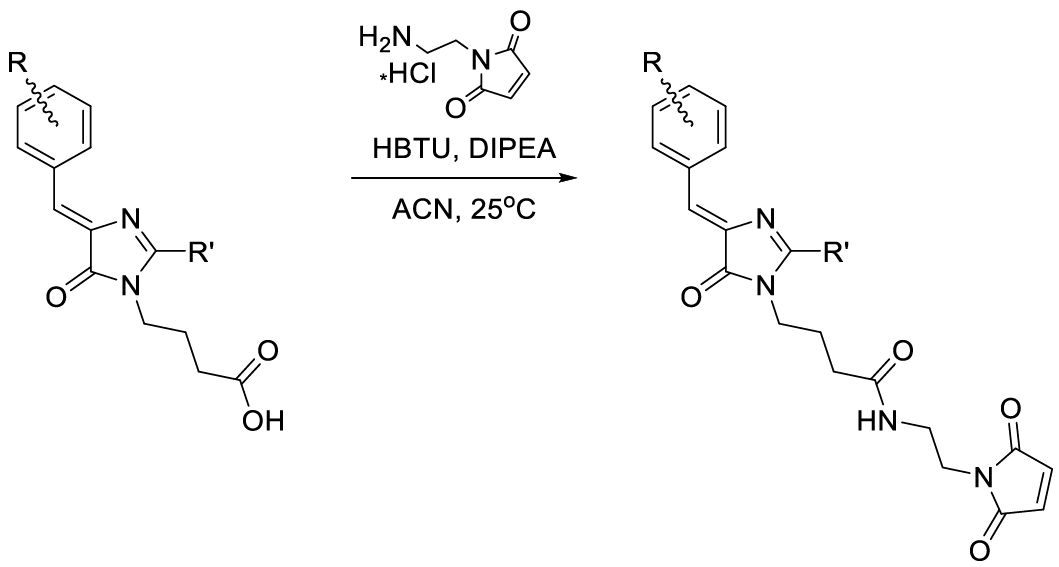

To the solution of 4-(4-((Z)-benzylidene)-2-((E)-styryl)-5-oxo-4,5-dihydro-1H-imidazol-1-yl)butanoic acids (0.1 mmol) in acetonitrile (2 mL) 1-(2-aminoethyl)-1H-pyrrole-2,5-dione hydrochloride (20 mg, 0.114 mmol), HBTU (40 mg, 0.106 mmol) and DIPEA (0.075 mL, 0.4 mmol) were added. The mixture was stirred for 12 h at room temperature. The mixture was dissolved with CHCl_3_ (200 mL), washed with a saturated solution of NaHCO_3_ (6 × 50), HCl (1%, 30 mL) and brain (2 × 50 mL) and dried over Na_2_SO_4_. The solvent was evaporated and the product was purified by column chromatography (CH_2_Cl_2_-MeOH, 100/5).

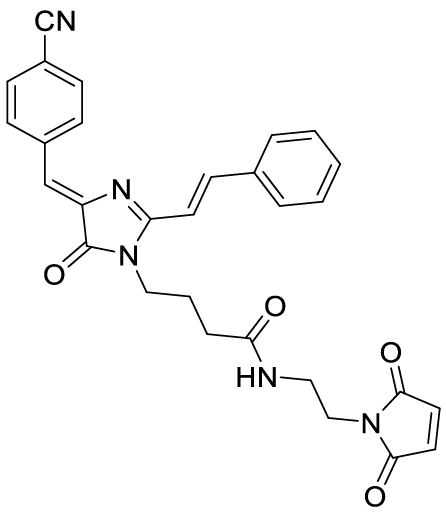

**4-(4-((*Z*)-4-cyanobenzylidene)-2-((*E*)-styryl)-5-oxo-4,5-dihydro-1*H*-imidazol-1-yl)-*N*-(2-(2,5-dioxo-2,5-dihydro-1*H*-pyrrol-1-yl)butanamide (DyeA maleimide)**

Yellow solid (38 mg, 75%); mp = 153-157 ºС; ^1^H NMR (700 MHz, DMSO-d_6_) δ ppm 8.48 (d, J=8.4 Hz, 2 H), 8.14 (d, J=15.6 Hz, 1 H), 7.98 (t, J=5.9 Hz, 1 H), 7.90 - 7.94 (m, 4 H), 7.46 - 7.53 (m, 3 H), 7.36 (d, J=15.8 Hz, 1 H), 7.08 (s, 1 H), 6.94 (s, 2 H), 3.76 (t, J=7.2 Hz, 2 H), 3.45 (t, J=5.8 Hz, 2 H), 3.19 (q, J=5.9 Hz, 2 H), 2.08 (t, J=7.2 Hz, 2 H), 1.77 (quin, J=7.2 Hz, 2 H); ^13^C NMR (176 MHz, DMSO-d_6_) δ ppm 171.6, 171.0, 169.8, 162.2, 142.0, 141.6, 139.0, 134.9, 134.4, 132.3, 132.3, 130.5, 129.0, 128.7, 122.0, 118.8, 113.5, 111.2, 38.2, 37.1, 36.9, 31.7, 24.8; HRMS (ESI) m/z: 508.1989 found (calcd for C_29_H_26_N_5_O_4_^+^, [M+H]^+^ 508.1979).

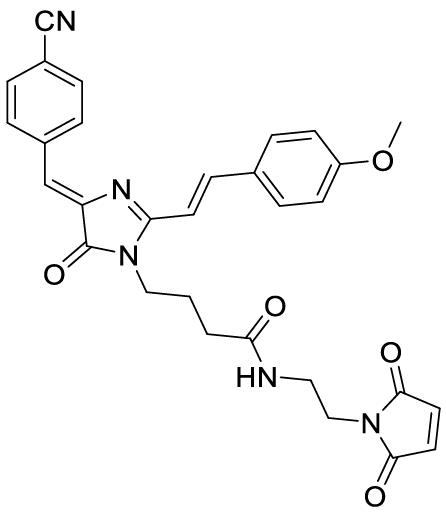

**4-(4-((*Z*)-4-cyanobenzylidene)-2-((*E*)-4-methoxystyryl)-5-oxo-4,5-dihydro-1*H*-imidazol-1-yl)-N-(2-(2,5-dioxo-2,5-dihydro-1H-pyrrol-1-yl)ethyl)butanamide (DyeB maleimide)**

Orange solid (33 mg, 61%); mp = 181-184ºС; ^1^H NMR and ^13^C NMR – see Supplementary Figure S11 bellow; HRMS (ESI) m/z: 538.2089 found (calculated for C_30_H_28_N_5_O_5_^+^, [M+H]^+^ 538.2085).

The structure of this compound was analyzed by two-dimensional NMR spectroscopy. The compound is represented in solution by two isomers in approximately 1:1 ratio. However, these isomers easily passed into each other in solution and the ratio could change with time. To elucidate the structure of both isomers, we performed full chemical shift assignment based on the HSQC, COSY, ^13^C- and ^15^N-HMBC spectra and measured the heteronuclear vicinal H-C J-couplings, using the PIP-HSQCMBC experiment [[1]](https://paperpile.com/c/v0RulZ/1z47).

According to the NMR data, this compound is present in solution as a mixture of Z and E isomers across the C1'-C2' double bond. The major state corresponds to the Z-isomer, which is supported by the large magnitude of H1'-H2' ^3^J-coupling (15.6 Hz), corresponding to the trans orientation of protons. In the minor state, the same J-coupling equals 13.0 Hz, which corresponds to the cis orientation. The maximal chemical shift difference between the two states is observed for the double bond protons H1' and H2', and exceeds 0.8 ppm for both of them.

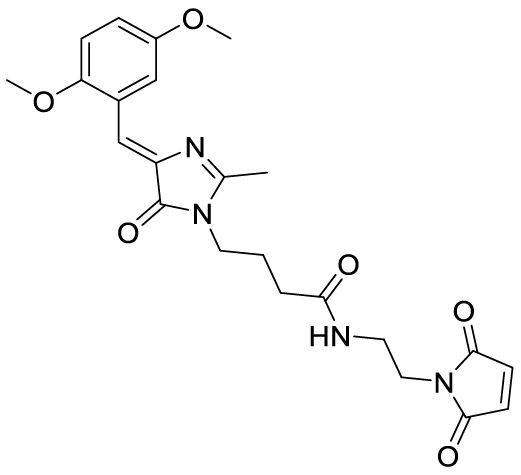

**(*Z*)-4-(4-(2,5-dimethoxybenzylidene)-2-methyl-5-oxo-4,5-dihydro-1*H*-imidazol-1-yl)-*N*-(2-(2,5-dioxo-2,5-dihydro-1*H*-pyrrol-1-yl)ethyl)butanamide (DyeC maleimide)**

Yellow solid (24 mg, 53%); mp = 125-128 ºС with decomposition; ^1^H NMR and ^13^C NMR see see Supplementary Figure S12 bellow; HRMS (ESI) m/z: 455.1915 found (calculated for C_23_H_27_N_4_O_6_^+^, [M+H]^+^ 455.1925).

The structure of this compound was analyzed by two-dimensional NMR spectroscopy. The compound is represented in solution by two isomers in approximately 1:2 ration (E and Z). However, these isomers easily passed into each other in solution and the ratio could change with time. To elucidate the structure of both isomers, we performed full chemical shift assignment based on the HSQC, COSY, ^13^C- and ^15^N-HMBC spectra and measured the heteronuclear vicinal H-C J-couplings, using the PIP-HSQCMBC experiment [[1]](https://paperpile.com/c/v0RulZ/1z47).

DyeC maleimide exists as two Z-E isomers across the C6-C4 double bond. The major form is the Z-isomer, which follows directly from the value of C6H-C5 J-coupling (4.8 Hz), which is indicative of the cis arrangement of the C6 proton and C5 carbon. In contrast, the corresponding J-coupling in the minor configuration of DyeC maleimide equaled 9.9 Hz, implying the trans arrangement of the proton and carbon and E-configuration of the double bond. This is supported by the chemical shift differences between the states, which are at most pronounced for the C6 carbon and exceed 10 ppm.

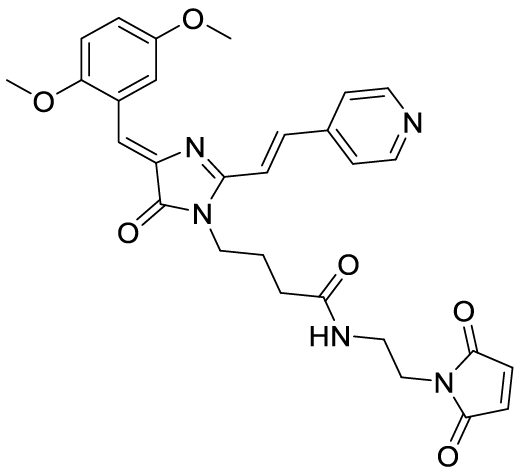

**4-(4-((*Z*)-2,5-dimethoxybenzylidene)-2-((*E*)-2-(pyridin-4-yl)vinyl)-5-oxo-4,5-dihydro-1*H*-imidazol-1-yl)-*N*-(2-(2,5-dioxo-2,5-dihydro-1*H*-pyrrol-1-yl)ethyl)butanamide (DyeD maleimide)**

Orange solid (22 mg, 41%); mp = 204-207ºС; ^1^H NMR (700 MHz, DMSO-d_6_) δ ppm 8.69 (d, J=6.1 Hz, 2 H), 8.53 (t, J=1.7 Hz, 1 H), 7.99 (t, J=6.2 Hz, 1 H), 7.92 (d, J=15.8 Hz, 1 H), 7.82 (d, J=5.9 Hz, 2 H), 7.59 (d, J=15.6 Hz, 1 H), 7.42 (s, 1 H), 7.05 (d, J=1.7 Hz, 2 H), 6.95 (s, 2 H), 3.86 (s, 3 H), 3.83 (s, 3 H), 3.75 (t, J=7.4 Hz, 2 H), 3.46 (t, J=5.8 Hz, 2 H), 3.20 (q, J=5.8 Hz, 2 H), 2.07 (t, J=7.3 Hz, 2 H), 1.76 (quin, J=7.3 Hz, 2 H); ^13^C NMR (201 MHz, DMSO-d_6_) δ ppm 171.7, 171.0, 169.7, 159.3, 153.4, 153.0, 150.3, 142.0, 138.7, 137.5, 134.4, 123.0, 122.1, 119.2, 118.5, 118.4, 116.2, 112.5, 56.2, 55.3, 39.0, 37.1, 36.9, 31.7, 24.9; HRMS (ESI) m/z: 544.2198 found (calcd for C_29_H_30_N_5_O_6_^+^, [M+H]^+^ 544.2191).

### Supplemental Figures

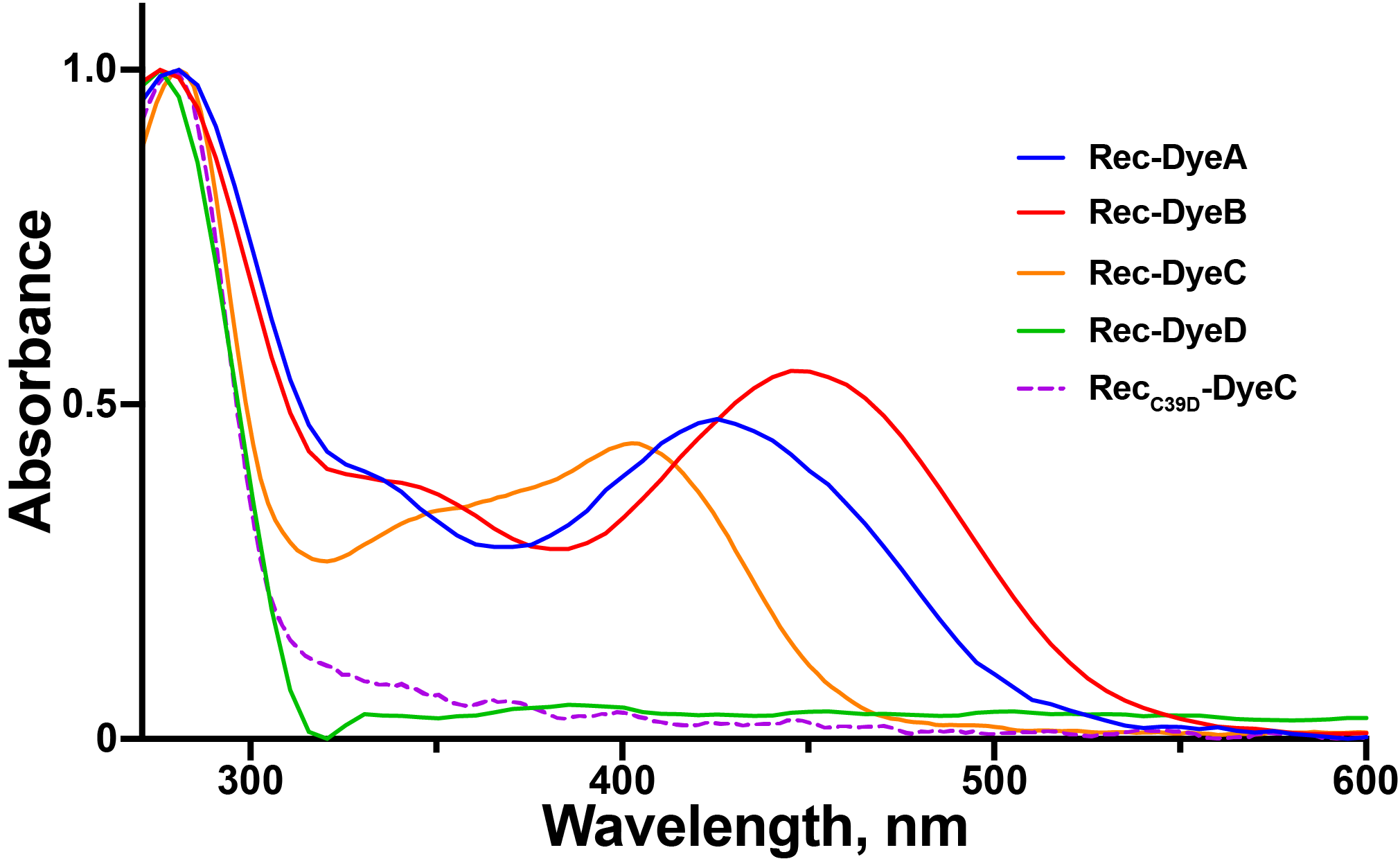

#### Supplementary Figure S1. Absorbance spectra of labeled Rec with all used dyes.

Rec_C39D_-DyeC is used as a control for labeling specificity. The absorption spectra are normalized at the protein absorption maximum (280 nm). Labeling efficiencies are ~ 90 %, 100 %, 85 %, and <5 % for Rec with DyeA, DyeB, DyeC, and DyeD, respectively, and <5 % for Rec_C39D_.

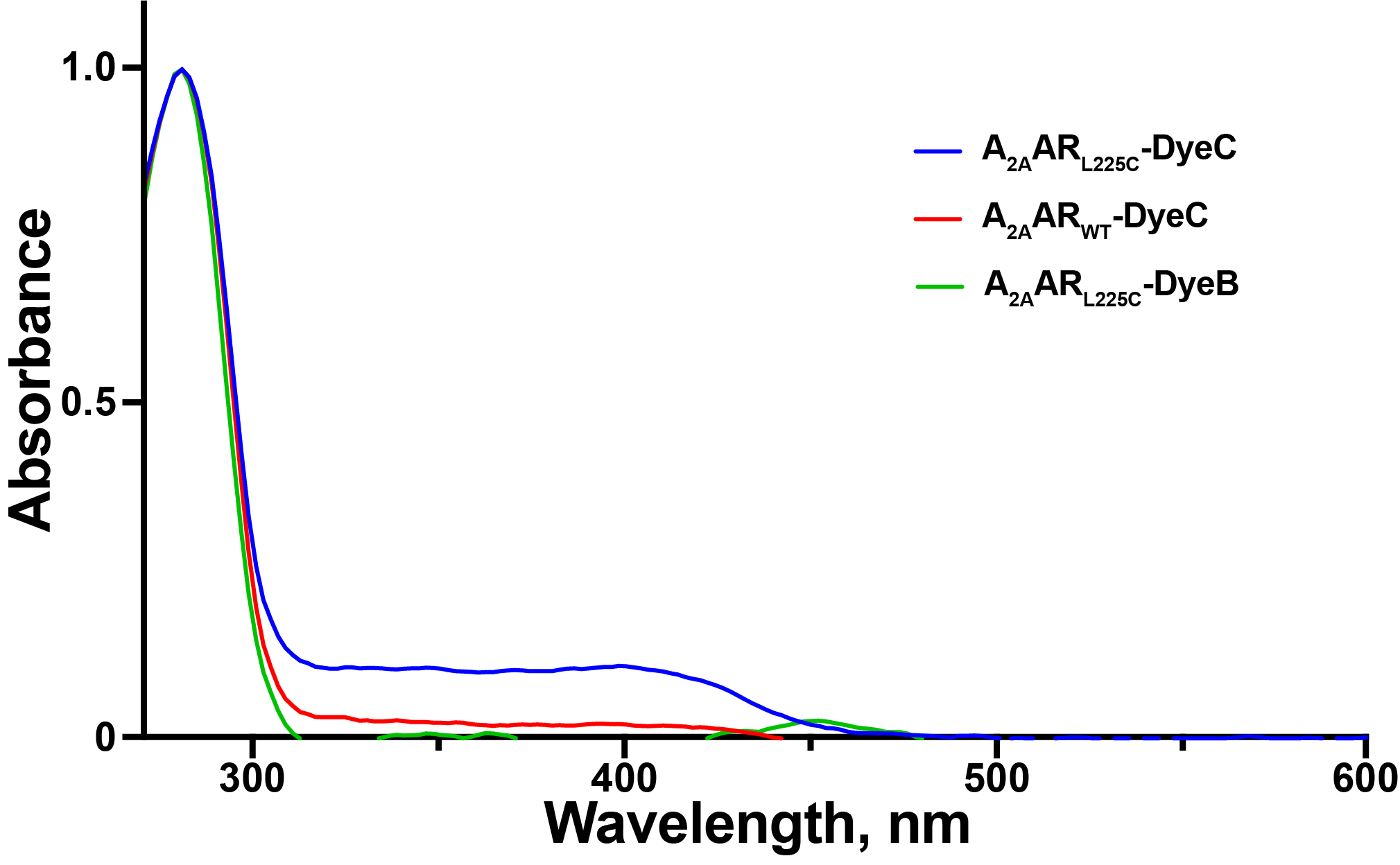

#### Supplementary Figure S2. Absorbance spectra of labeled A_2A_AR.

A_2A_AR_WT_-DyeC is used as a control for labeling specificity. The absorption spectra are normalized at the protein absorption maximum (280 nm). Labeling efficiencies are <5 % and ~ 80 % for A_2A_AR_L225C_ with dyes B and C, respectively, and <10 % for A_2A_AR_WT_-DyeC.

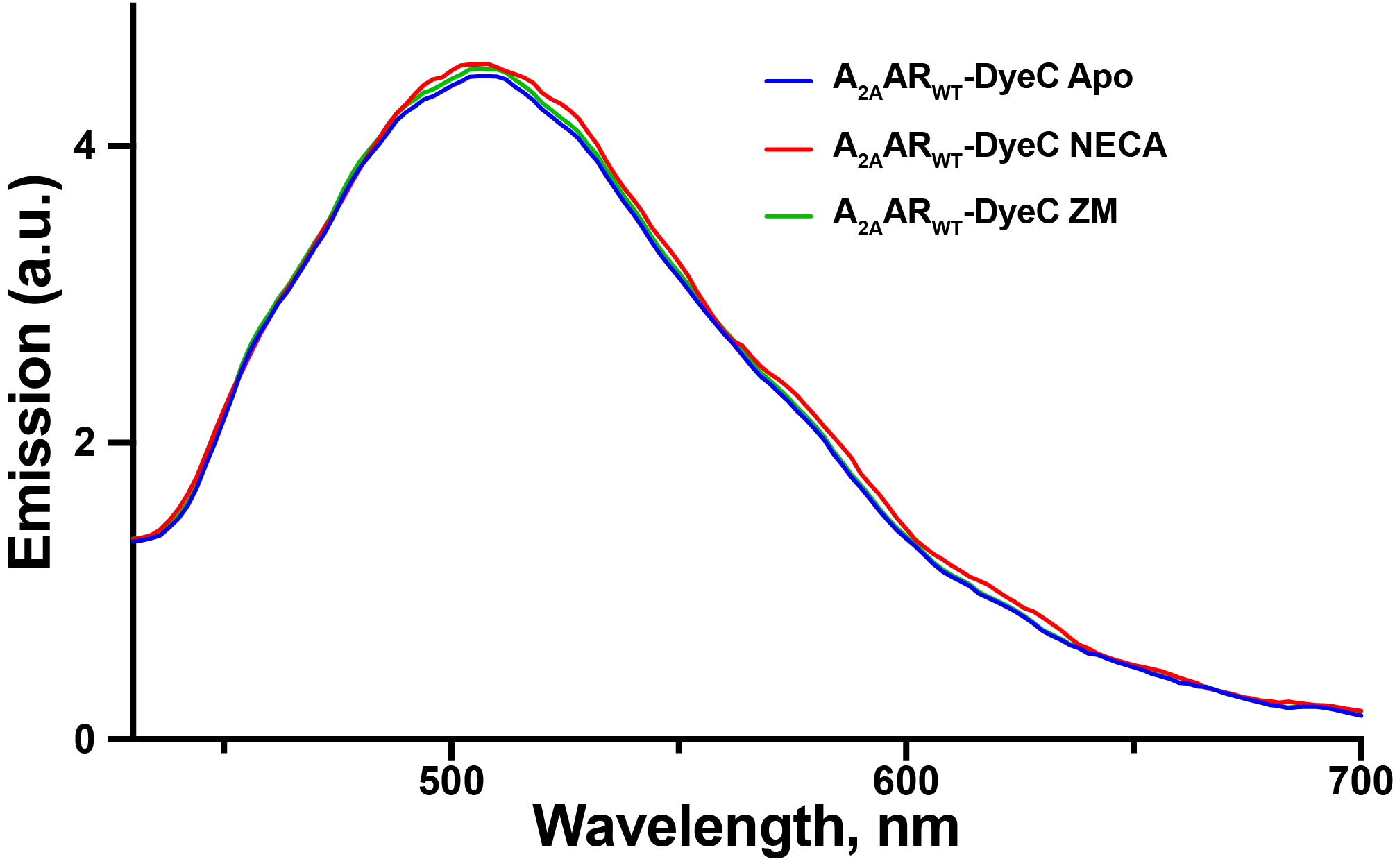

#### Supplementary Figure S3. Fluorescence emission spectra of A_2A_AR_WT_-DyeC with added ligands.

A_2A_AR_WT_-DyeC shows no response to the agonist NECA as opposed to A_2A_AR_L225C_-DyeC. This result suggests that there is no contribution to the change in A_2A_AR_L225C_-DyeC emission from nonspecific labeling. Excitation wavelength was 410 nm.

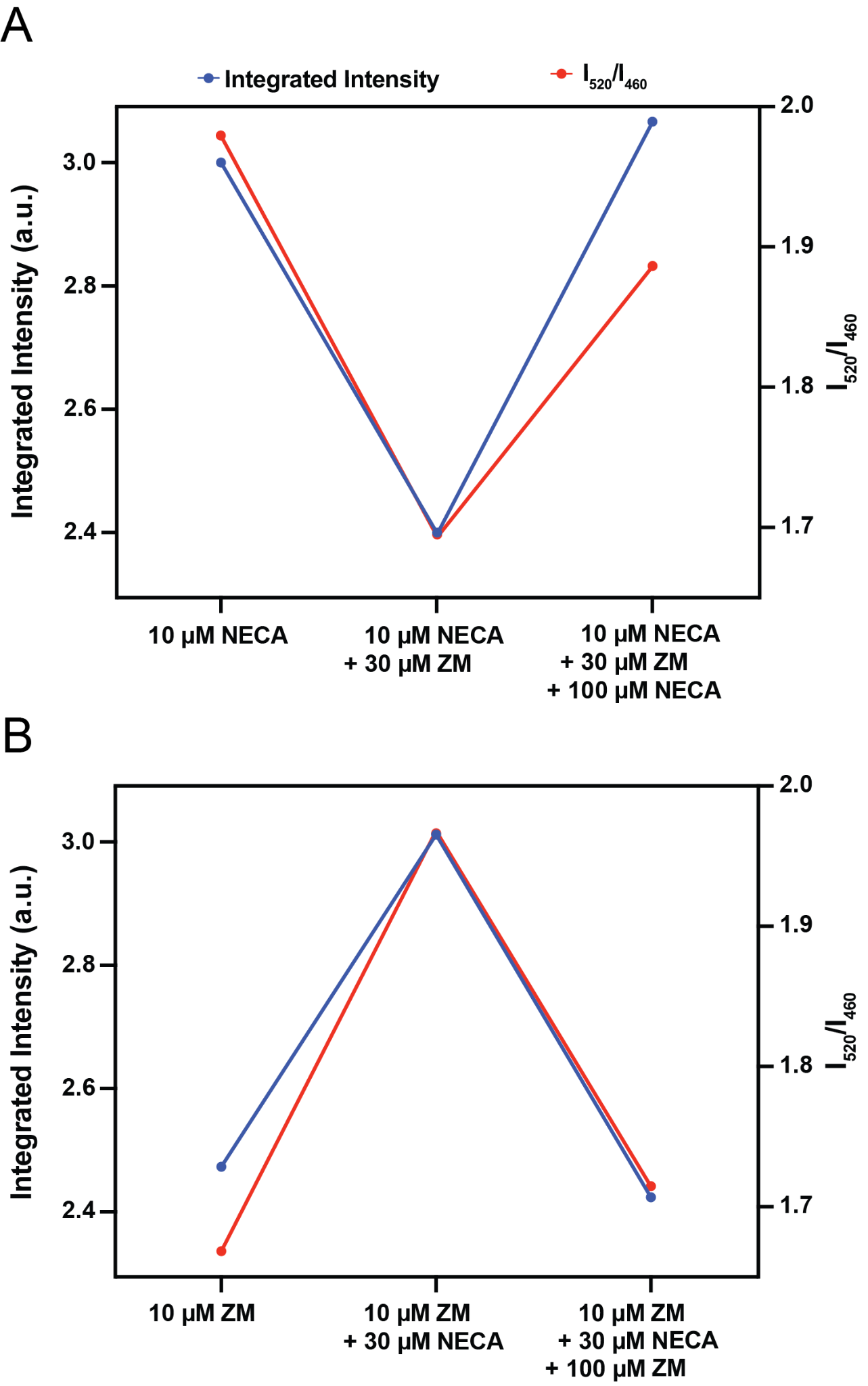

#### Supplementary Figure S4. Displacement of ligands in A_2A_AR_L225C_-DyeC.

Two experiments were performed. In the first experiment (**A**), 10 µM NECA was added initially, followed by the addition of 30 µM ZM241385, and finally NECA was added again to a final concentration of 110 µM. In the second experiment (**B**), 10 µM ZM241385 was added initially, followed by the addition of 30 µM NECA, and finally ZM241385 was added again to a final concentration of 110 µM. Protein concentration was 2 µM. Excitation wavelength was 410 nm.

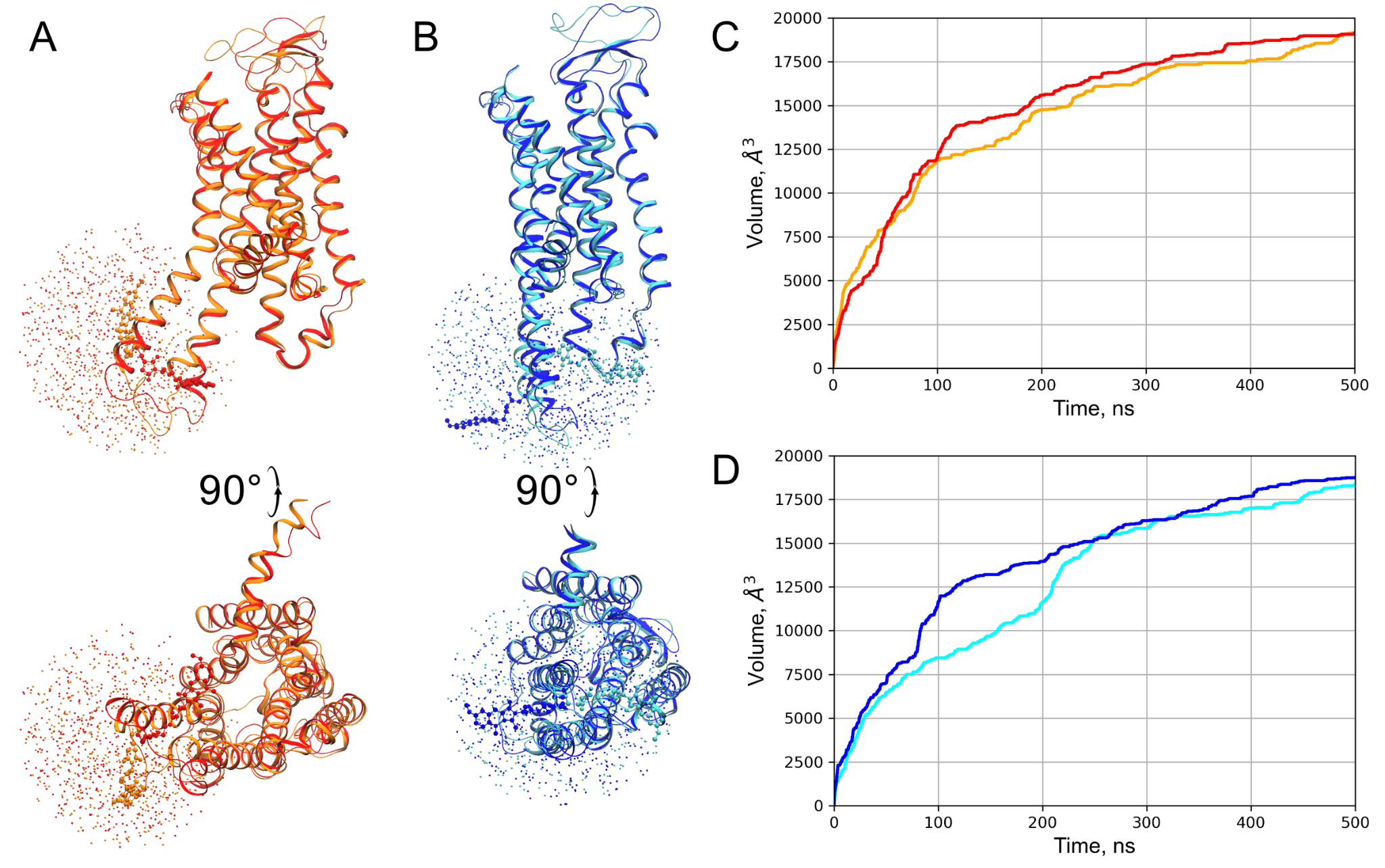

#### Supplementary Figure S5. Metadynamics convergence analysis.

**A-B:** Positions of the dimethoxybenzene ring of the DyeC label throughout the metadynamics simulations shown every 1 ns. Each simulation was run for 500 ns. Results for two replicate simulations are shown for each receptor state, in orange and red for the active state, in cyan and blue - for the inactive state. Lipids and water are not shown for clarity. **C-D:** The cumulative volume explored by the fluorescent label plotted as a function of the simulation time for two replicate simulations of the active (panel C) and inactive (panel D) receptor states, colors match those in panels A-B.

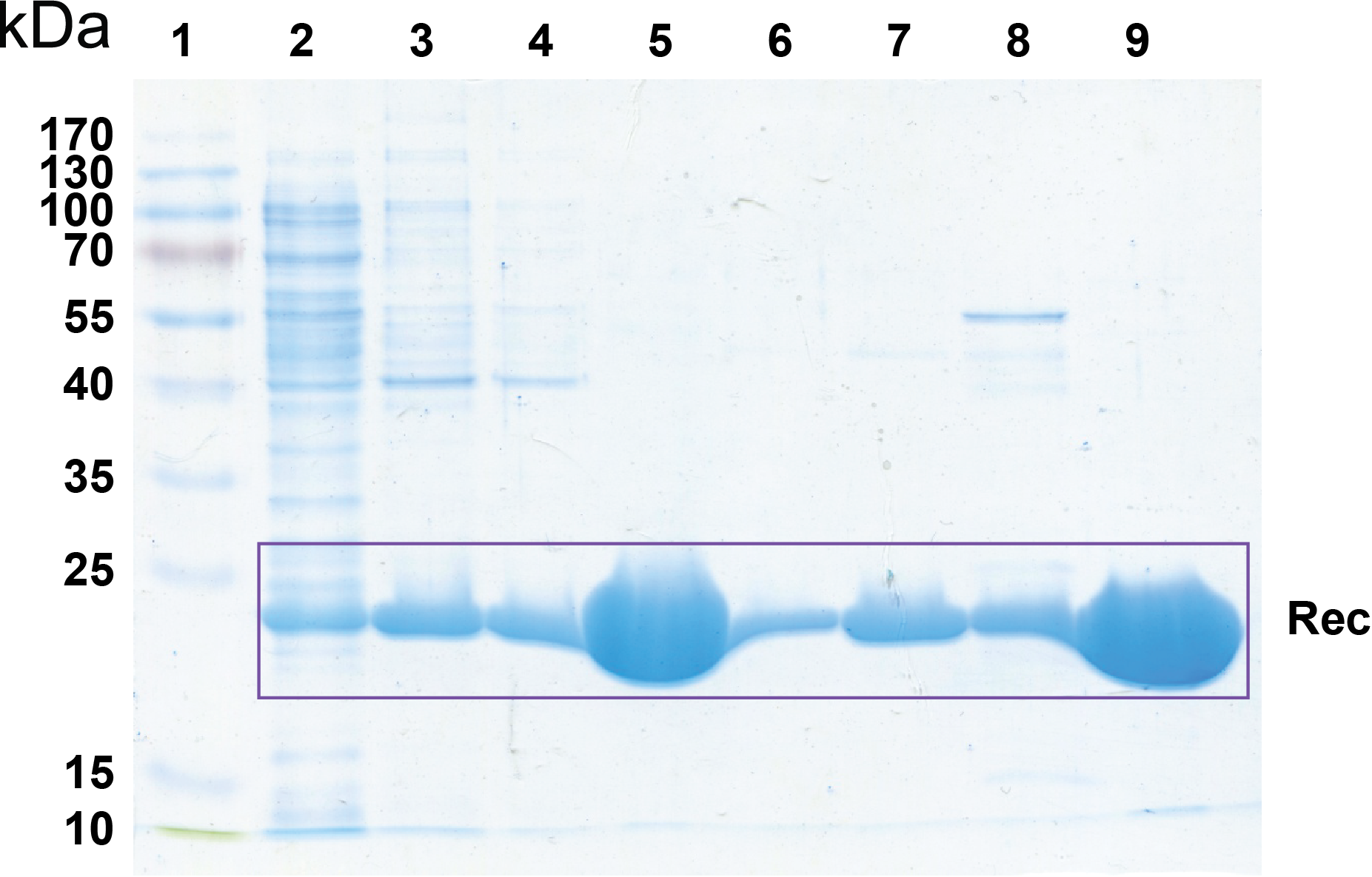

#### Supplementary Figure S6. SDS-PAGE (12.5%) of fractions obtained during purification of Rec/Rec_C39D_.

Track 1 – Thermo Scientific™ PageRuler™ Prestained Protein Ladder; Track 2 – Bacterial lysate (Escherichia coli strain BL21-CodonPlus®(DE3)-RIL-X (Agilent), pBB131, pET11d-Rec); Track 3 – Chromatography on Phenyl Sepharose (Cytiva): eluate in 20 mM Tris-HCl, pH=7.5, 1 mM EGTA, 2 mM MgCl_2_ and 1 mM DTT; Tracks 4-8 – Chromatography on HiTrap Q FF (Cytiva): fractions of gradient 0-1 M NaCl in 20 mM Tris-HCl, pH=7.5, 1 mM DTT; Track 5 – Selected fraction of myristoylated wild type Rec; Track 9 – Selected fraction of myristoylated Rec_C39D_, purified exactly as described for wild type Rec. The calculated Rec mass is 23.0 kDa.

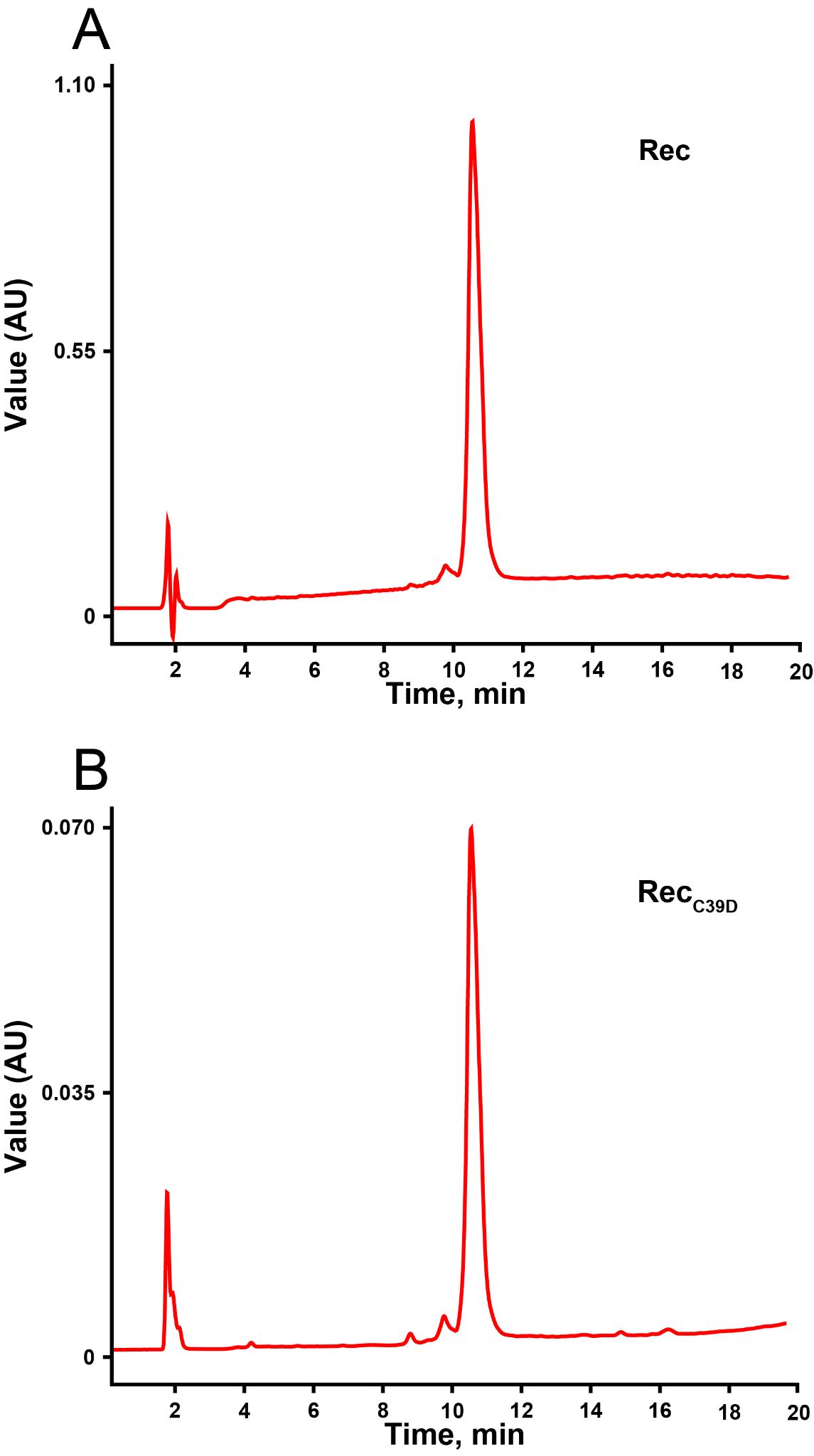

#### Supplementary Figure S7. Analytical HPLC of Rec and Rec_C39D_ in the acetonitrile-water system.

Chromatograms of purified Rec (A) and Rec_C39D_ (B) obtained using Breeze QS HPLC System (Waters) equipped with Luna C18 reversed-phase column (Phenomenex) in acetonitrile gradient. Retention time: non-myristoylated Rec/Rec_C39D_ – 9.7 min, myristoylated Rec/Rec_C39D_ – 10.7 min.

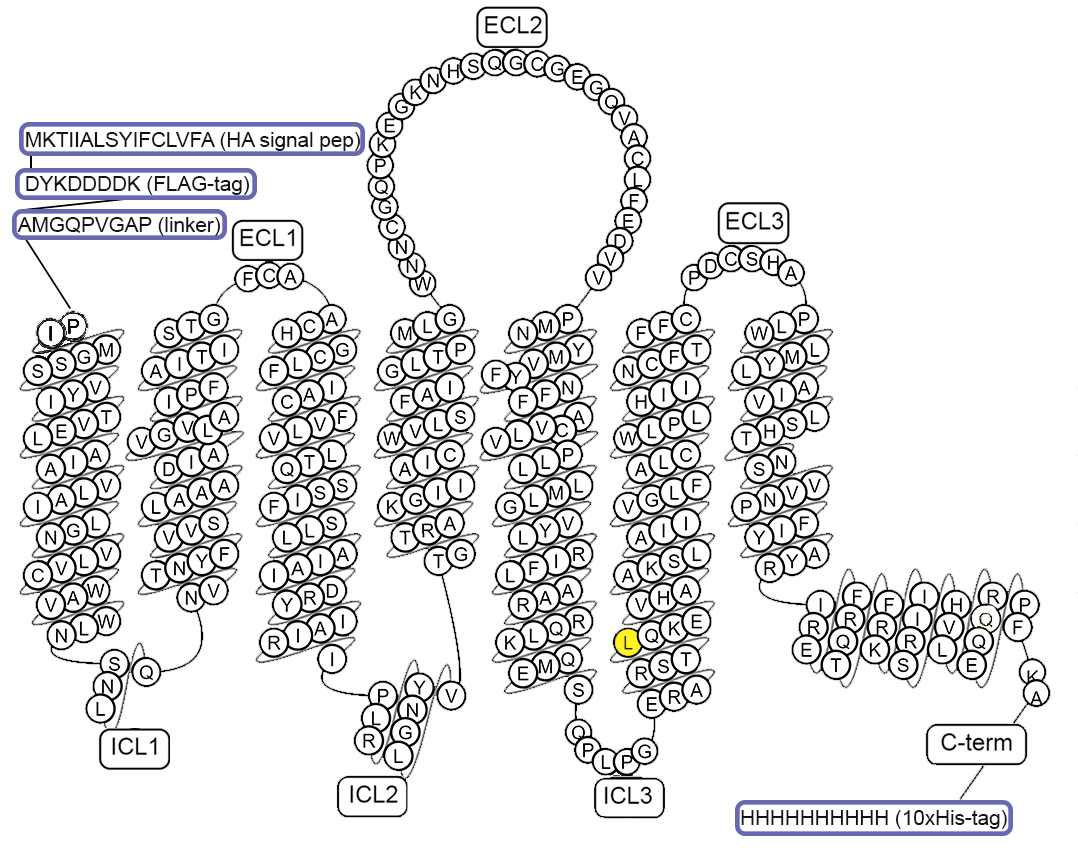

#### Supplementary Figure S8. A snake-plot diagram of the A_2A_AR construct used in this work.

L225^6.27^ residue mutated to cysteine for the labeling is colored in yellow. The snake-plot was drawn using the GPCRdb.org website.

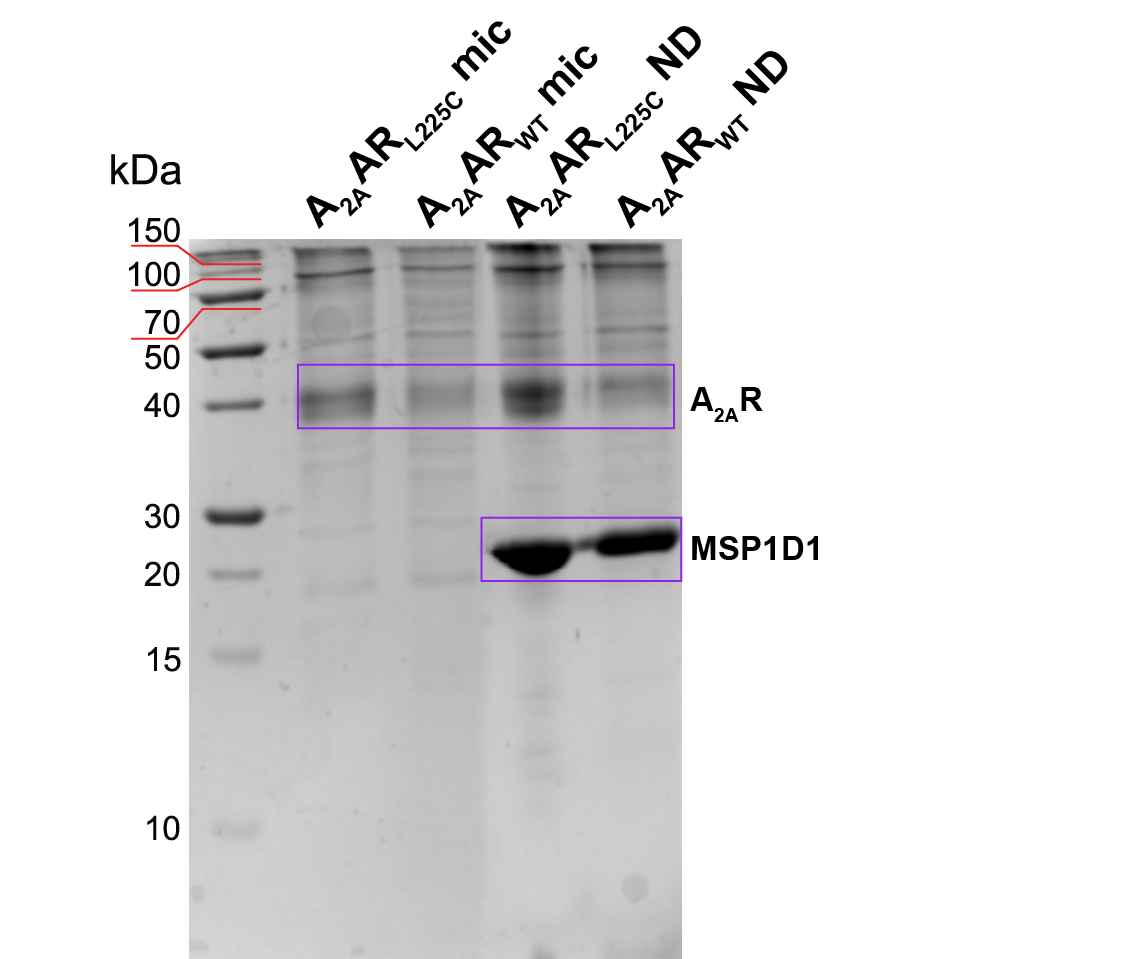

#### Supplementary Figure S9. SDS-PAGE of A_2A_AR-DyeC samples.

A_2A_AR_WT_-DyeC and A_2A_AR_L225C_-DyeC in micelles showed bands around 40 kDa (the calculated A_2A_AR mass is 39.8 kDa), A_2A_AR_WT_-DyeC and A_2A_AR_L225C_-DyeC reconstituted in ND showed two bands: for A_2A_AR around 40 kDa, and for MSP1D1 around 20 kDa (the calculated MSP1D1 mass is 24.7 kDa).

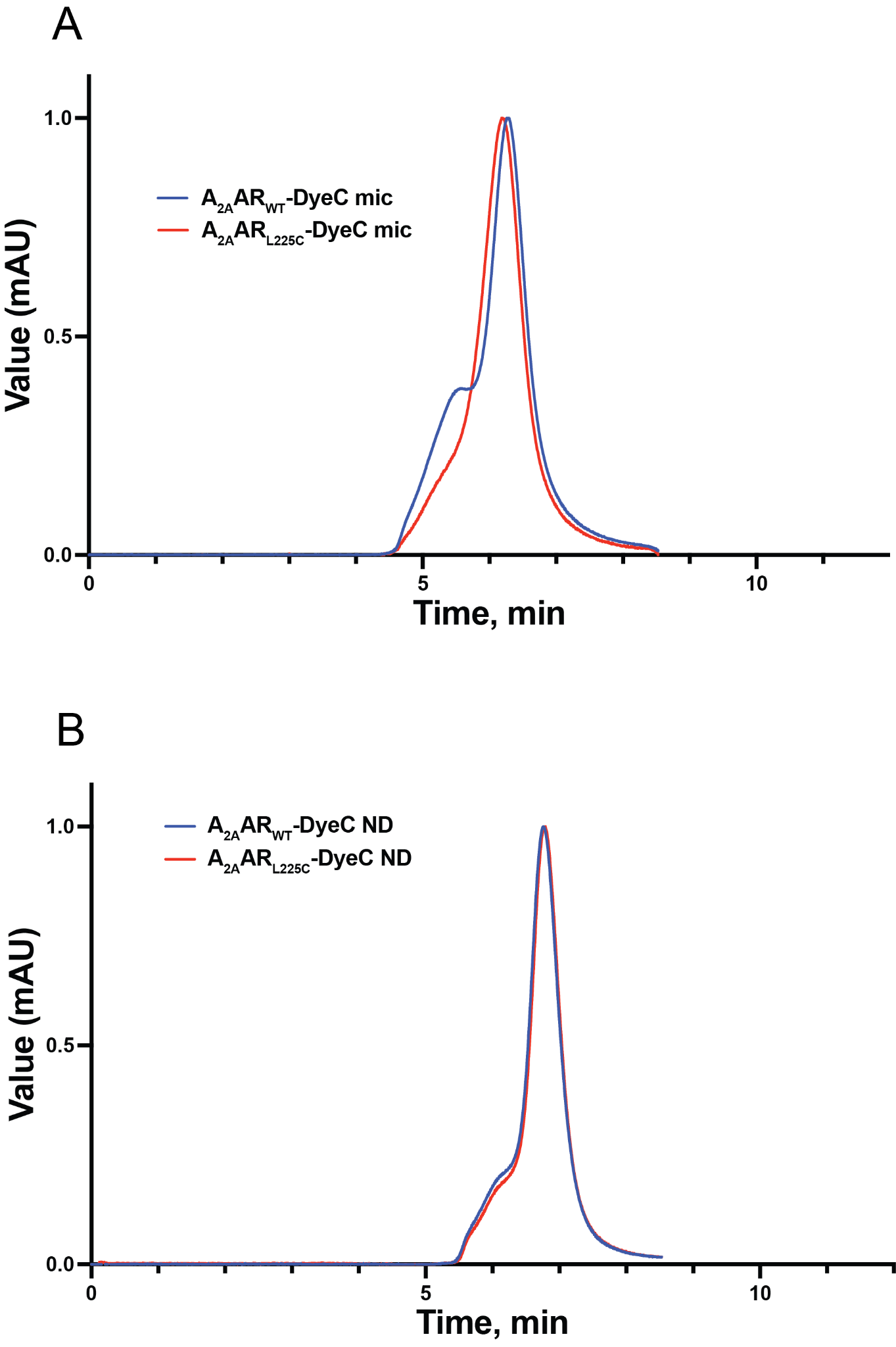

#### Supplementary Figure S10. SEC of A_2A_AR in micelles and nanodiscs.

A: Normalized analytical SEC of A_2A_AR_L225C_-DyeC and A_2A_AR_WT_-DyeC in micelles. B: Normalized analytical SEC of A_2A_AR_L225C_-DyeC and A_2A_AR_WT_-DyeC in ND. Chromatograms show mostly monomeric protein in both membrane-modeling systems.

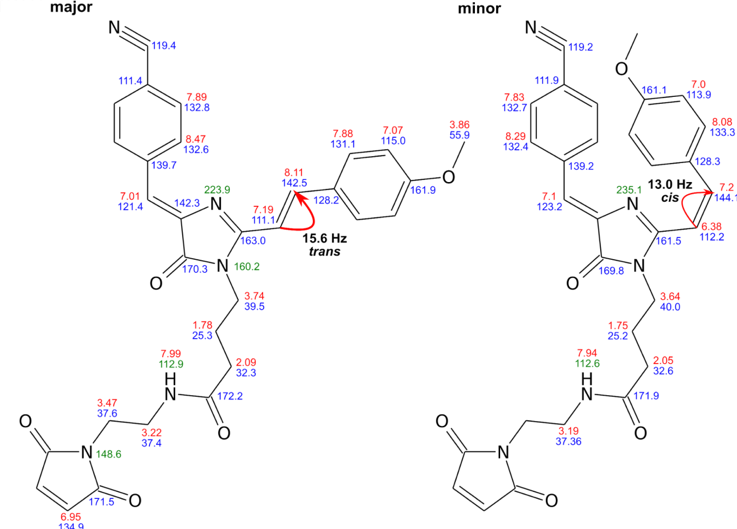

#### Supplementary Figure S11. Structures of DyeB maleimide isomers.

Chemical shift assignments are indicated by labels colored in red (^1^H), blue (^13^C), and green (^15^N). The red arrows denote the J-coupling between the double bond protons. Only the different chemical shifts in the minor state are indicated.

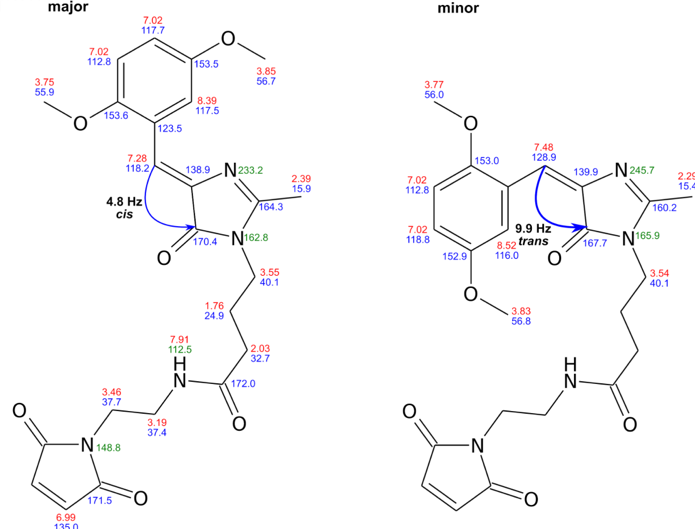

#### Supplementary Figure S12. Structures of DyeC maleimide isomers.

Chemical shift assignments are indicated by labels colored in red (^1^H), blue (^13^C) and green (^15^N). The blue arrows denote the J-coupling between the double bond proton and carbonyl carbon. Only the different chemical shifts in the minor state are indicated.

#### Supplementary Table Sr1. Solvatochromic properties of maleimide compounds DyeA, DyeB, DyeC, and DyeD.

|  |  |  |  | DyeA | | | DyeB | | | DyeC | | | DyeD | | |
| --- | --- | --- | --- | --- | --- | --- | --- | --- | --- | --- | --- | --- | --- | --- | --- |
|  | Solvent | logP^*^ | Viscosity, mPa⋅s^*^ | λ_ABS_, nm | λ_EM_, nm | FQY, % | λ_ABS_, nm | λ_EM_, nm | FQY, % | λ_ABS_, nm | λ_EM_, nm | FQY, % | λ_ABS_, nm | λ_EM_, nm | FQY, % |
| 1 | Water | -1.38 | 0.9 | 416 | 557 | 0.4 | 434 | 575 | 0.69 | ~393 | ~570 | 0.6 | ~450 | ~643 | N/D |
| 2 | DMSO | -1.35 | 2.47 | 431 | 568 | 1.9 | 444 | 585 | 2.16 | 399 | 502 | 17.8 | 461 | 590 | 4.1 |
| 3 | DMF | -1.01 | 0.8 | 428 | 563 | 1.7 | 443 | 580 | 1.45 | 395 | 498 | 14.1 | 458 | 581 | 6.2 |
| 4 | MeOH | -0.77 | 0.54 | 421 | 559 | 1.1 | 436 | 575 | 0.7 | 398 | 539 | 2.9 | 457 | 600 | 0.4 |
| 5 | ACN | -0.34 | 0.35 | 424 | 564 | 1.5 | 437 | 585 | 0.82 | 394 | 502 | 7.3 | 452 | 583 | 4.4 |
| 6 | EtOH | -0.31 | 1.07 | 424 | 555 | 1.4 | 440 | 575 | 1.11 | 398 | 522 | 4 | 459 | 592 | 1.2 |
| 7 | Dioxane | -0.27 | 1.2 | 425 | 554 | 3.4 | 441 | 570 | 1.88 | 394 | 477 | 13.4 | 452 | 564 | 22.4 |
| 8 | Acetone | -0.24 | 0.32 | 425 | 563 | 1.7 | 437 | 580 | 0.13 | 393 | 485 | 12.2 | 450 | 576 | 9.2 |
| 9 | PrOH-1 | 0.25 | 2.26 | 424 | 559 | 1.7 | N/M | N/M | N/M | 398 | 517 | 3.8 | 460 | 588 | 2 |
| 10 | THF | 0.46 | 0.53 | 427 | 560 | 2.3 | 442 | 580 | 1.5 | 393 | 475 | 10.2 | 451 | 566 | 19.8 |
| 11 | PY | 0.65 | 0.89 | 431 | 565 | 2.6 | 446 | 585 | 2.43 | 398 | 489 | 11.2 | 463 | 582 | 11 |
| 12 | EtOAc | 0.73 | 0.42 | 425 | 554 | 2.3 | 438 | 575 | 1.37 | 392 | 476 | 10.1 | 452 | 564 | 18.7 |
| 13 | BuOH | 0.88 | 2.57 | N/M | N/M | N/M | 442 | 570 | 1.66 | N/M | N/M | N/M | N/M | N/M | N/M |
| 14 | Et_2_O | 0.89 | 0.24 | 424 | 554 | 2.6 | 440 | 570 | 1.33 | 392 | 466 | 9.5 | 446 | 560 | 19.4 |
| 15 | CH_2_Cl_2_ | 1.25 | 0.44 | 425 | 560 | 2.2 | 441 | 580 | 1.28 | 397 | 495 | 4 | 457 | 579 | 7.9 |
| 16 | Pentanol-1 | 1.51 | 3.5 | N/M | N/M | N/M | N/M | N/M | N/M | 401 | 507 | 4.9 | N/M | N/M | N/M |
| 17 | Toluene | 2.73 | 0.56 | 429 | 557 | 3.8 | 443 | 575 | 1.97 | 396 | 475 | 9.7 | 456 | 543 | 21.4 |
| 18 | Decanol-1 | 4.57 | 10.9 | N/M | N/M | N/M | N/M | N/M | N/M | 401 | 491 | 9.4 | N/M | N/M | N/M |
| 19 | Undecanol-1 | 4.72 | 17.2 | N/M | N/M | N/M | 446 | 570 | 2.92 | 402 | 486 | 10.5 | N/M | N/M | N/M |
| 20 | Octane | 5.18 | 0.52 | 426 | 550 | 2.3 | 442 | 565 | 1.5 | 396 | 458 | 5.7 | 449 | 560 | 4.1 |
| 21 | Dodecane | 6.1 | 1.36 | ~428 | ~556 | N/D | 439 | 565 | 2.28 | 396 | 458 | 6.3 | ~451 | ~567 | N/D |
| 22 | Pentadecane | 7.71 | 1.95 | N/M | N/M | N/M | 443 | 565 | 2.53 | 398 | 462 | 7.9 | N/M | N/M | N/M |

^*^ – data from pubchem.ncbi.nlm.nih.gov/compound

N/D – not determined due to low fluorescence

N/M – not measured

#### Supplementary Table S2. Integrated Intensity and I_520_/I_460_ ratios for A_2A_AR_L225C_-DyeC with various ligands.

| **Ligand** | **I_520_/I_460_** | **Integrated Intensity** |
| --- | --- | --- |
| Apo | 1.76 ± 0.05 (n = 11) | 2.64 ± 0.06 (n = 11) |
| ZM241385 | 1.68 ± 0.04 (n = 9) | 2.62 ± 0.07 (n = 9) |
| SCH58261 | 1.66 ± 0.04 (n = 9) | 2.66 ± 0.08 (n = 9) |
| Adenosine | 2.06 ± 0.03 (n = 10) | 3.15 ± 0.08 (n = 10) |
| NECA | 1.99 ± 0.05 (n = 15) | 3.16 ± 0.03 (n = 15) |
| HMA | 1.42 ± 0.03 (n = 9) | 3.03 ± 0.04 (n = 9) |

Mean ± SD are given for a sample of repeated experiments. For each condition, protein from at least three different protein purifications was used, total numbers of technical repeats are given in brackets.

#### Supplementary Table S3. Detection of GPCR structural changes with environmentally sensitive dyes.

| **Dye** | **Protein** | **Labelling position** | **Additional quencher** | **Mutation** | **Lipid Modelling System** | **Max. intensity change, %** | **Max. λ_EM_ shift, nm** | **λ_EM_, nm** | **λ_EXC_, nm** | **ε, M^-1^⋅cm^-1^⋅10^3^** | **Ref.** |
| --- | --- | --- | --- | --- | --- | --- | --- | --- | --- | --- | --- |
| IANBD | β_2_AR | undefined native Cys | - | - | DDM | 5 | 0 | 523 | 481 | 21 [[2–4]](https://paperpile.com/c/v0RulZ/rPRix+vps2g+9zq9j) | [[2–4]](https://paperpile.com/c/v0RulZ/rPRix+vps2g+9zq9j) |
| FM | β_2_AR | C265^6.27^ | - | - | DDM | 15 | N/A | 520 | 490 | 83 [[5]](https://paperpile.com/c/v0RulZ/gpZRJ) | [[5,6]](https://paperpile.com/c/v0RulZ/gpZRJ+4nqjX) |
| FM | β_2_AR | C265^6.27^ | K224-oxyl-NHS | All K to R/Q224^5.63^K | DDM | ~23 | N/A | 520 | 490 | 83 [[5]](https://paperpile.com/c/v0RulZ/gpZRJ) | [[5]](https://paperpile.com/c/v0RulZ/gpZRJ) |
| TMR-5 | β_2_AR | C265^6.27^ | - | - | DDM | ~20–~60 | N/A | 571 | 541 | 101 [[7]](https://paperpile.com/c/v0RulZ/lr7oO) | [[8–10]](https://paperpile.com/c/v0RulZ/uotuX+thKAM+A9GmC) |
| APM | β_2_AR | C265^6.27^ or C271^6.33^ | - | C378/406S or C378/406S, C265^6.27^A | DDM | 0 | <2 | 625 | 515 | ~50 [[11]](https://paperpile.com/c/v0RulZ/EH42S) | [[11]](https://paperpile.com/c/v0RulZ/EH42S) |
| Bimane | β_2_AR | C271^6.33^ | I135W | C77^2.48^V, C265^6.27^/378/406A C327^7.54^S | DDM+CHEMS | ~50 | 0 | 458 | 370 | 5 [[12]](https://paperpile.com/c/v0RulZ/Bjqzf) | [[13]](https://paperpile.com/c/v0RulZ/9uObN) |
| Bimane | β_2_AR | C265^6.27^ | - | 365-trun. or T4 lysozyme fusion | DDM | <26 | <8 | 448 | 350 | 5 [[12]](https://paperpile.com/c/v0RulZ/Bjqzf) | [[14]](https://paperpile.com/c/v0RulZ/loupv) |
| Bimane | β_2_AR | C265^6.27^ | - | C77^2.48^V, C327^7.54^S, C378/406A | ND | 50 | 15 | 450 | 370 | 5 [[12]](https://paperpile.com/c/v0RulZ/Bjqzf) | [[15]](https://paperpile.com/c/v0RulZ/yOHqv) |
| TMR-5 | β_2_AR | C265^6.27^ | - | C77^2.48^V, C327^7.54^S, C378/406A | DDM | 15 | 4 | 550 | 515 | 101 [[7]](https://paperpile.com/c/v0RulZ/lr7oO) | [[16]](https://paperpile.com/c/v0RulZ/N5K4X) |
| Bimane | GHSR | C304^7.34^ or C146^3.55^ | - | C146^3.55^S or C304^7.34^S | ND | ~55* | ~20 | ~450 | 375 | 5 [[12]](https://paperpile.com/c/v0RulZ/Bjqzf) | [[17]](https://paperpile.com/c/v0RulZ/FXkkf) |
| Bimane | Rhodopsin | C141^3.56^ or C150^4.39^ or C228^5.63^ or C258^6.41^ or C250^6.33^ | KI | C140^3.55^/316^8.53^/322^8.59^/323S and K141^3.56^C or E150^4.39^C or F228^5.63^C or V258^6.41^C or V250^6.33^C | DDM | ~50 | ~100 | 460 | 380 | 5 [[12]](https://paperpile.com/c/v0RulZ/Bjqzf) | [[18]](https://paperpile.com/c/v0RulZ/rDxbz) |
| Bimane | Rhodopsin | C227^5.62^ or C244^6.27^ or C250^6.33^ or C251^6.34^ | KI | C140^3.55^/316^8.53^/322^8.59^/323S and V227^5.62^C or Q244^6.27^C or V258^6.41^C or V250^6.33^C | DDM | N/A | 12 | ~460 | 380 | 5 [[12]](https://paperpile.com/c/v0RulZ/Bjqzf) | [[19]](https://paperpile.com/c/v0RulZ/ziM1Q) |
| Bimane | Parapinopsin | C227^5.62^ or C244^6.27^ or C250^6.33^ or C251^6.34^ | KI | V138^3.53^F, C140A, C316^6.40^A, C323A and M227^5.62^C or A244^6.27^C or M258^6.40^C or V250^6.33^C | DDM | N/A | 7 | ~465 | 380 | 5 [[12]](https://paperpile.com/c/v0RulZ/Bjqzf) | [[19]](https://paperpile.com/c/v0RulZ/ziM1Q) |
| IAEDANS | A_2A_AR | C231^6.33^ | - | A231^6.33^C | SMALP | 50** | 0 | 460 | 340 | 6 [[20]](https://paperpile.com/c/v0RulZ/ax4lC) | [[21]](https://paperpile.com/c/v0RulZ/Ne23V) |
| DyeC | A_2A_AR | C225^6.27^ | - | L225^6.27^C, 316-trun | ND | 20 | 5 | 515 | 410 | 14 | present study |

* - opposite to other studies, the maximum effect was caused by inverse agonist but not agonist.

** - opposite to other studies, the maximum effect was caused by antagonist but not agonist.

N/A - not available
